# Proteomic analysis of SUMO1-SUMOylome changes during defense elicitation in *Arabidopsis*

**DOI:** 10.1101/2020.08.02.233544

**Authors:** Kishor D. Ingole, Shraddha K. Dahale, Saikat Bhattacharjee

**Affiliations:** Laboratory of Signal Transduction and Plant Resistance, UNESCO-Regional Centre for Biotechnology (RCB), NCR Biotech Science Cluster, 3^rd^ Milestone, Faridabad-Gurgaon Expressway, Faridabad- 121 001, Haryana, India; Kalinga Institute of Industrial Technology (KIIT) University, Bhubaneswar- 751 024, Odisha, India

**Keywords:** Post-translational modification (PTM), SUMOylation, autoimmunity, *PstDC3000*, LC-MS/MS, plant-pathogen interaction, plant defense response

## Abstract

Rapid adaptation of plants to developmental or physiological cues is facilitated by specific receptors that transduce the signals mostly via post-translational modification (PTM) cascades of downstream partners. Reversible covalent attachment of SMALL UBIQUITIN-LIKE MODIFIER (SUMO), a process termed as SUMOylation, influence growth, development and adaptation of plants to various stresses. Strong regulatory mechanisms maintain the steady-state SUMOylome and mutants with SUMOylation disturbances display mis-primed immunity often with growth consequences. Identity of the SUMO-substrates undergoing SUMOylation changes during defences however remain largely unknown. Here we exploit either the auto-immune property of an Arabidopsis mutant or defense responses induced in wild-type plants against *Pseudomonas syringae* pv *tomato* (*PstDC3000*) to enrich and identify SUMO1-substrates. Our results demonstrate massive enhancement of SUMO1-conjugates due to increased SUMOylation efficiencies during defense responses. Of the 261 proteins we identify, 29 have been previously implicated in immune-associated processes. Role of others expand to diverse cellular roles indicating massive readjustments the SUMOylome alterations may cause during induction of immunity. Overall, our study highlights the complexities of a plant immune network and identifies multiple SUMO-substrates that may orchestrate the signalling.

## INTRODUCTION

Post-translational modifications (PTMs) by reversible attachment of the SMALL UBIQUITIN-LIKE MODIFIER (SUMO), through a process termed as SUMOylation, dynamically and reversibly modulate target protein fate, localization, or function (reviewed in Morrell and Sadanandom, 2019). In plants, abiotic or biotic cues cause massive changes on global SUMOylome (Bailey et al. 2015; Miller et al. 2010; Colignon et al. 2017a). Though primarily utilized to regulate nuclear functions such as DNA replication, chromatin remodeling, transcription in eukaryotes, role of SUMOylation extend to other organellar processes (Colignon et al. 2017a; Miller et al. 2010). Nevertheless, the vast majority of nuclear proteins as SUMO-substrates is indicative of the rapid mechanism to modulate transcriptome in response to stimulus.

SUMOylation cascade recruits processed SUMOs to covalently attach via an isopeptide bond to ε-amino group of lysine residues on target proteins. The modified lysine often is a part of a partially conserved motif, ψ-K-X-D/E (ψ = hydrophobic amino acid, X= any amino acid, D/E= aspartate/glutamate). A conjugatable SUMO, subsequently forms a thiol-ester bond with the SUMO E1 ACTIVATING ENZYME (SAE). A trans-esterification reaction further shuttles SUMO to the SUMO E2 CONJUGATING ENZYME 1 (SCE1) and then to target lysine on SUMOylation substrates. SUMO1 can conjugate to itself and form poly-SUMO1 chains (Miller et al. 2010). SUMO-substrate specificities are regulated by SUMO E3 LIGASES such as HIGH PLOIDY2/METHYL METHANE SULFONATE21 (HPY2/MMS21) and SAP and MIZ1 (SIZ1) (Morell and Sadanandom, 2019). Fate of covalently-attached SUMOs may either include de-conjugation and recycling by SUMO proteases such as EARLY IN SHORT DAYS 4 (ESD4), its closest homolog ESD4-LIKE SUMO PROTEASE 1 (ELS1), or targeted proteolysis of the substrate through ubiquitin-mediated pathway. SUMO-interactions are also non-covalent in nature facilitated by hydrophobic amino acid-rich SUMO-interaction motifs (SIMs) present in the recipient. SIMs populate several SUMOylation-associated proteins implying strong auto-regulatory mechanisms (Wang et al. 2010). Unlike fruit fly, worm, or yeast, humans and plants express multiple SUMO isoforms (Flotho and Melchior, 2013). In *Arabidopsis thaliana*, 4 SUMO isoforms are expressed namely SUMO1, SUMO2, SUMO3 and SUMO5 suggestive of substrate specificities and/or SUMO crosstalks (van den Burg et al. 2010).

Distinctions between self and non-self in plants is facilitated by the robust two-layered immune system (Jones and Dangl, 2006). Primary defense signalling routes responds to recognition via surface-localized pattern-recognition receptors (PRRs) for conserved molecular signatures called pathogen- or microbe-associated molecular patterns (PAMPs/MAPMs) present on pathogens, to elicit PAMP-triggered immunity (PTI). Manipulative pathogens secrete effectors to suppress PTI which in turn are detected by intracellular resistance (R) proteins to activate a more amplified effector-triggered immunity (ETI). ETI in many cases lead to a programmed cell death called hypersensitive response (HR) (Jones and Dangl, 2006). The hormone salicylic acid (SA) plays a central role in plant immunity and its accumulation drives intricately interconnected networks causing increased expression of *PATHOGENESIS RELATED PROTEIN 1/2 (PR1/2*) and other defense-associated genes (Loake and Grant, 2007). A key mediator of SA-signaling is ENHANCED DISEASE SUSCEPTIBILITY 1 (EDS1) the null-mutant of which dramatically decreases plant combat abilities (Wiermer et al. 2005). Preventing mis-priming of immunity and regulating of its amplitudes especially in the context of activation of R proteins is performed by negative regulators such as SUPPRESSOR OF rps4-RLD1 (SRFR1) (Kim et al. 2010). A *srfr1-4* mutant is growth retarded, has elevated SA-based defences, and is enhanced resistant to the bacterial pathogen *Pseudomonas syringae* pv *tomato DC3000 (PstDC3000*). *SRFR1* encodes a tetratricopeptide (TPR) protein with versatile interactors such as multiple R proteins, EDS1, co-chaperone SUPPRESSOR OF THE G2 ALLELE OF skp1 (SGT1), and members of the TEOSINTE BRANCHED1/CYCLOIDEA/PCF (TCP) transcription factors (Bhattacharjee et al. 2011; Kim et al. 2014, 2010; Li et al. 2010). Loss of *EDS1* abolishes growth retardation and increased immunity of *srfr1-4* implicating constitutive SA-signaling sectors for these defects. These features make *srfr1-4* an excellent system for studying defense signaling processes.

Host SUMOylome readjustments influence immune outcomes and been comprehensively highlighted in several excellent reviews (Augustine and Vierstra, 2018; van den Burg and Takken, 2010). Arabidopsis *siz1-2*, SUMO protease mutants *OVERLY TOLERANT TO SALT 1/2 (OTS1/2*), or *esd4-1* plants accumulate elevated SA, constitutively express PR proteins and are increased resistant to *PstDC3000* (Lee *et al*. 2006; Bailey *et al*. 2015; Villajuana-Bonequi *et al*. 2014). Plants null for *SUM1* and expressing microRNA-silenced *SUM2 (sum1-1 amiR-SUM2*) also have heightened immunity (van den Burg et al. 2010). Constitutive activation of SA-dependent defenses occur in *SUM1/2/3*-overexpressing transgenic plants (van den Burg et al. 2010). From these reports, it is increasingly evident that unregulated increases (*eds4-1, ots1/2* or over-expressing *SUM1/2*) or decreases (in *siz1-2* or *sum1-1 amiR-SUM2*) in global SUMOylome bear immune consequences. It is also not surprising that pathogens attempt to manipulate host SUMOylome to increase their colonization efficiencies (Wimmer & Schreiner, 2015; Verma et al. 2018). Several bacterial phytopathogenic effectors interfere with host SUMOylation as a mode to suppress immunity (Hotson et al. 2003). XopD, a secreted effector from *Xanthomonas campestris pv. vesicatoria* (*Xcv*) de-conjugates SUMO from unknown targets in plants (Tan et al. 2015). Mutations that disrupt SUMO-protease functions of XopD not only render the cognate strain deficient in virulence but also lower defense induction in the host plant.

Focused approaches to identify basal or differential pool of SUMOylated proteins in response to stress exposures have gained increasing prominence in plant systems (Elrouby and Coupland, 2010; López-Torrejón et al. 2013; Miller et al. 2010; Park et al. 2011). A major breakthrough in the enrichment methodologies was reported by Miller et al. (2010). Because of a longer native SUMO1-footprint causing difficulties in detection of modifications via mass spectrometric methods, a tagged SUMO1 variant (His-H89R-SUMO1) was developed that functionally complemented the defects of loss of *SUM1 in planta*. Stringent purification protocols minimizing false-positives identified a plethora of targets ascribed to multiple biological pathways. Perhaps more elegantly, SAE2, SCE1, SIZ1, and EDS4 were also identified as candidates whose SUMOylation footprints changed upon stress exposure implying that SUMOs themselves moderate SUMOylation proficiencies in cellular responses. More recently, a modified three-dimensional gel-electrophoresis (3D SDS-PAGE)-based protocol involving differential labeling of proteins with fluorescent dyes was developed to identify SUMOylated candidates differing between susceptible or resistant cultivars of potato to *Phytophthora infestans* infections (Colignon et al. 2017b). Remarkably, the authors observed that resistance was associated with the ability of the host to rapidly achieve SUMO-conjugate increments whereas the susceptible cultivar was deficient in this process and succumbed to manipulative efforts of the pathogen.

In this study, we developed, exploited, and compared two independent systems to identify changes in SUMOylated proteome in context of defense responses. In each of these systems, the expression of the tagged SUMO1-variant His-H89R-SUMO1 (Miller et al. 2010) facilitated rigid enrichment and identification of SUMO-substrates via liquid chromatography followed by mass spectrometry analysis (LC-MS/MS). The auto-immune *srfr1-4* system exhibited massive increase in basal SUMO1/2-conjugates relative to the wild-type plants and thus provided a good source to enrich for SUMO-candidates. Wild-type plants challenged with *PstDC3000* showed progressive increases in SUMO1/2-conjugate accumulations and therefore served as a physiological pathosystem for identifying SUMOylated proteins in defense responses. Overall, the two systems shared 57.9% common SUMO-substrates encompassing diverse GO categories. Our findings provide deeper dimensions into cellular networks and targets impinged by SUMOylome changes thus reflecting the adjustments of a plant during immunity.

## RESULTS

### Arabidopsis auto-immune mutant *srfr1-4* show net enhancements in global SUMO1/2-SUMOylome

The auto-immune *srfr1-4* plants are developmentally stunted, have elevated SA levels accompanied by constitutive upregulation of PR1/PR2 (Kim et al. 2010) (Figure 1A,B). The complemented line expressing HA-epitope-tagged genomic *SRFR1* sequences from its native promoter (*srfr1-4:HA-SRFR1g*) restores *srfr1-4* defects and have been described earlier (Kim et al. 2010). The auto-immune feature of *srfr1-4* makes it an excellent system to explore immune-associated changes. Similar systems have been explored earlier to identify key immune modulators and decipher the complexities of defense signaling networks (van Wersch et al. 2016). To explore SUMOylome changes associated with immunity we probed total extracts from wild-type (Col-0), *srfr1-4*, and *srfr1-4:HA-SRFR1g* plants with anti-SUMO1/2 antibodies. Because of strong identity between SUMO1 and SUMO2 isoforms, the antibodies cross-reacts with both proteins (van den Burg et al. 2010). Marked enhancement in global SUMO1/2-SUMOylome and free-SUMO1/2 proteins (~12 kDa) is strikingly apparent in the *srfr1-4* extracts with clear restoration to Col-0 levels in the complemented line (Figure 1C). Introducing SA-signaling deficiencies through the *EDS1* mutation (*eds1-2*) in *srfr1-4* abolishes its growth defects and auto-immunity (Bhattacharjee et al. 2011). Remarkably, in these *srfr1-4 eds1-2* plants, increased SUMO1/2-conjugates are brought down to Col-0 levels implicating SA-regulated and EDS1-dependent perturbations for SUMOylome upregulations in *srfr1-4* (Figure S1).

**Figure 1.**
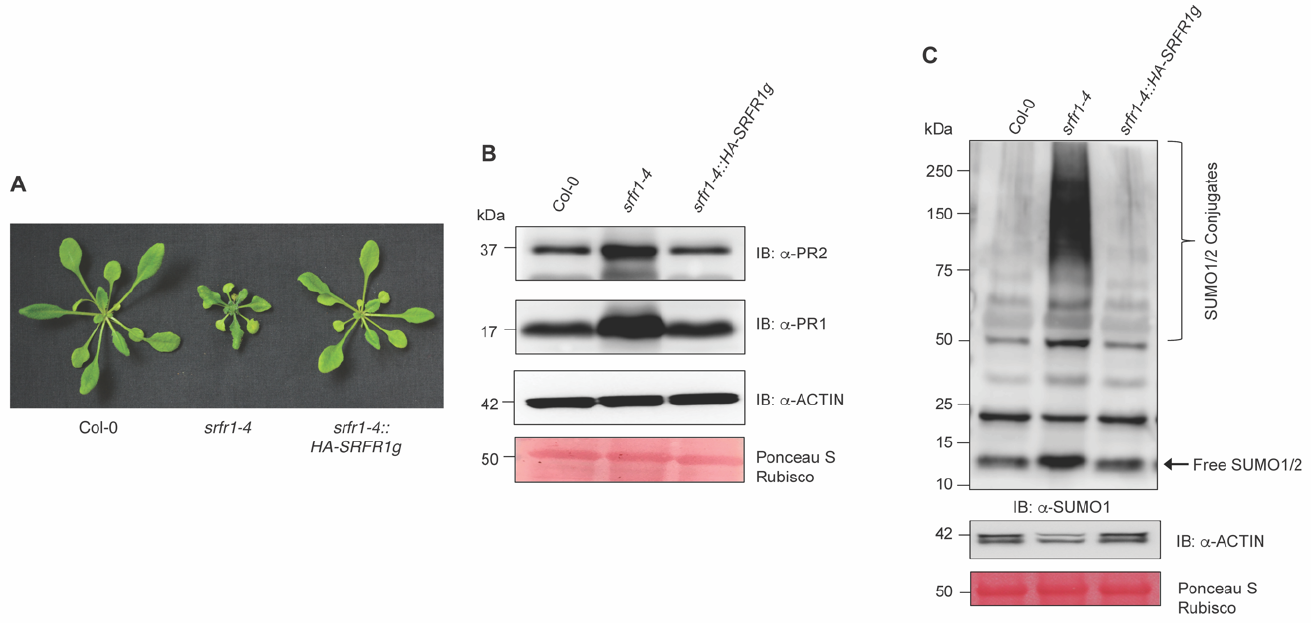
Enhanced net SUMO1/2-SUMOylome and free-SUMO1/2 levels in the auto-immune *srfr1-4* plants. **(A)** Developmental phenotype of 3-weeks-old Col-0, *srfr1-4* and *srfr1-4::HA-SRFR1g* plants. **(B)** Basal levels of defense markers PR1 and PR2 are elevated in *srfr1-4*. **(C)** Increased accumulation of SUMO1/2-conjugates and free-SUMO1/2 in *srfr1-4*. Total protein extracts of indicated plants were resolved in a 10% (B) or 4-15% gradient (C) SDS-PAGE gels and immunoblots (IB) performed with indicated antibodies. Actin immunoblots and PonceauS staining of the PVDF membranes indicate comparable protein loading across samples. Migration position of molecular weight standards (in kDa) are marked at left of the respective blots. Positions of SUMO1/2-conjugates and free-SUMO1/2 are shown.

### *SUM1* transcripts undergo increased translation during immunity

Escalated SUMO1/2-conjugates in *srfr1-4* may be attributed but not limited to increased expression/activity of SUMO1/2, SUMO E1 (SCE1), E2 (SAE2) or E3 (SIZ1, HPY2) enzymes, or decreased functions of SUMO deconjugases (EDS4, ELS1) influenced transcriptionally or post-transcriptionally. Although SUMO1/2-conjugates and free-SUMO1/2 are augmented upon SA-application, *SUM3*, but not *SUM1/2* transcriptions are SA-inducible (van den Burg et al. 2010). In *srfr1-4*, we note similar elevations in *SUM3* expressions although *SUM1/2* transcript levels remain Col-0-comparable (Figure 2A). Relative transcript levels of *SAE1a*, *SAE2*, *SCE1*, *SIZ1* and *HPY2* are interestingly increased whereas *ESD4* and *ELS1* are comparatively lower in *srfr1-4* than Col-0 (Figure 2B-D). These results suggest that SUMOylation efficiencies are more promoted in *srfr1-4*. As the levels of free-SUMO1/2 are higher in *srfr1-4*, we tested whether existing *SUM1/2* transcripts are more translated. Polysomes contain ribosome-loaded mRNAs undergoing active translation (Chassé et al. 2017)(Figure 2E). Polysome-associated transcripts in *srfr1-4* are ~2.5-fold more enriched for *SUM1*, but not *SUM2* mRNAs, than Col-0 or the complemented line (Figure 2F). Transcripts of *PR1*, as expected and in accordance to their increased protein abundance are also considerably higher in these fractions from *srfr1-4*. Taken together, heightened SUMOylation efficiencies, in part also contributed by more availability of SUMO1 proteins, are responsible for global enhancement of SUMO1/2-conjugates in *srfr1-4*.

**Figure 2.**
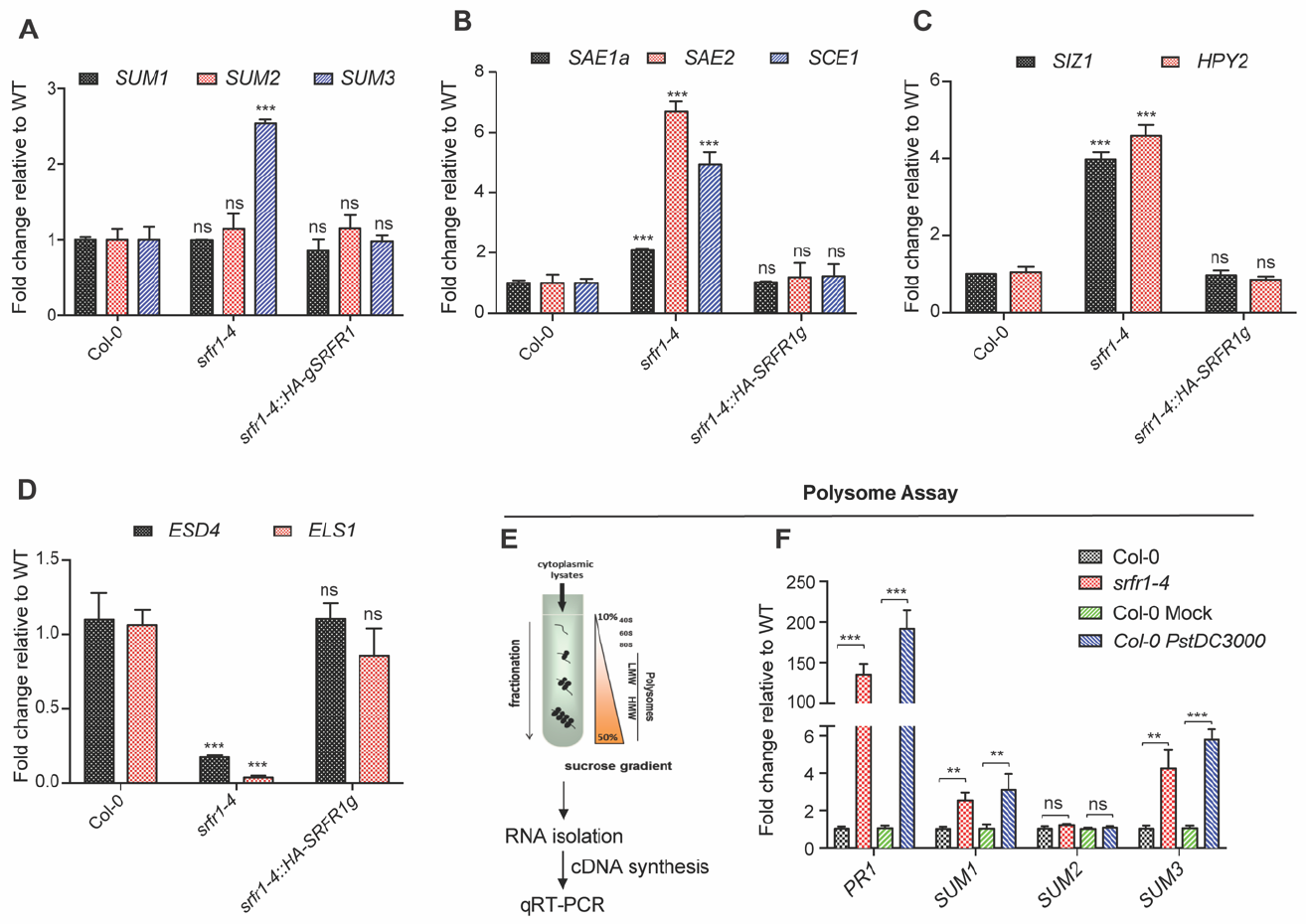
Increased SUMO1-conjugation efficiencies in *srfr1-4*. **(A-D)** Relative expression levels of SUMO isoform *SUM1, SUM2, SUM3* (A), SUMO E1, E2 enzymes *SAE1a, SAE2, SCE1* (B), SUMO E3 ligases *SIZ1, HPY2* (C), SUMO de-conjugases *ESD4, ELS1* (D) genes in 3-weeks-old Col-0, *srfr1-4* and *srfr1-4::HA-SRFR1g* plants. **(E)** Flow chart and principle of polysome-enrichments for qRT-PCRs. **(F)** Relative enrichments of *PR1*, *SUM1*, *SUM2*, and *SUM3* transcripts in polysomes isolated from Col-0, *srfr1-4*, Col-0 Mock, and Col-0 *PstDC3000*-infected samples. Data normalization was performed to *MON1* (At2g28390) expression. The values are represented as fold-change relative to Col-0 (WT). Data is representative of mean of three biological replicates (n=3). Error bars indicate SD. Student *t*-test was performed to calculate statistical significance; **=*p*<0.01; ***=*p*<0.001, ns= not significant.

### SUMO1/2-SUMOylome is kinetically induced upon *PstDC3000* infections

Changes in global SUMOylome has not been reported during *PstDC3000* challenges. To investigate this, Col-0 plants were infiltrated with the bacterial suspension, total protein extracts prepared at progressive time points post-infection, and analyzed for SUMO1/2-SUMOylome changes. Distinct accelerated increase in SUMO1/2-conjugates is detected till 24-hpi (hrs post-infection) with moderate reduction at 48-hpi (Figure 3A). Expression of *PR1* shows a similar trend likely suggesting a link between SA-signaling route and SUMOylome changes (Figure 3B). And as is noted for SA-treatments (van den Burg et al. 2010), *PstDC3000* infections also cause time-dependent increase in *SUM3* expressions reaching plateau at 24-hpi and toning-down by 48-hpi (Figure 3C). Expressions of *SUM1* or *SUM2* genes remain unaffected during these time-points. Although *SAE1a* or *HPY2* transcript levels are not affected during the infection stages, *SAE2, SCE1*, and *SIZ1* expressions increase gradually (Figure 3D,E). We especially note that *ESD4* and *ELS1* expressions are dramatically suppressed as early as 3-hpi with progressive increases at later stages of infection reaching basal levels by 48-hpi. Similar to *srfr1-4, SUM1, PR1, SUM3*, but not *SUM2*, mRNAs presence in polysomes is enhanced in 24-hpi Col-0 *PstDC3000*-infected samples than the mock control (Figure 2F). Overall, these results lead us to conclude that net enhancements in SUMO1/2-SUMOylome during defense induction are orchestrated primarily by improved SUMOylation efficiencies.

**Figure 3.**
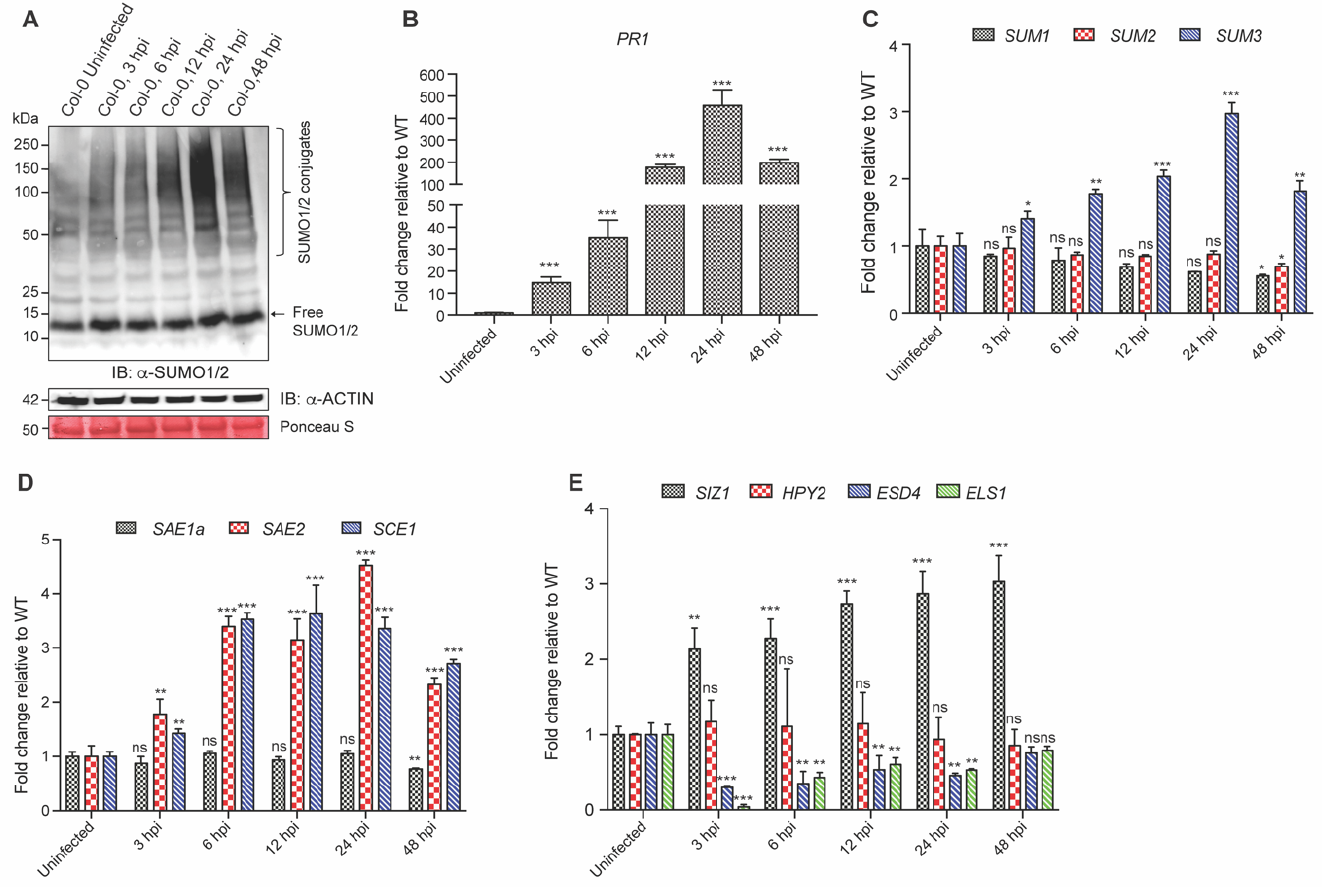
SUMO1/2-conjugations are promoted during *PstDC3000*-challenges. **(A)** SUMO1 immunoblot on total protein extracts from Col-0 uninfected and *PstDC3000*-infected tissues at indicated time-points (hpi: hours post-infection). Total proteins were resolved in a 4-15% gradient SDS-PAGE gel and immunoblotted with anti-SUMO1/2 antibodies. Positions of SUMO1/2-conjugates and free-SUMO1/2 are indicated. Actin immunoblot and PonceauS staining of Rubisco are indicative of comparable protein loadings. Relative migrations of molecular weight standards (in kDa) are indicated. **(B-E)** Relative expression kinetics of *PR1* (B), *SUM1, SUM2, SUM3* (B), *SAE1a, SAE2, SCE1* (C), *SIZ1, HPY2, ESD4, ELS1* (D) in uninfected, or in *PstDC3000*-infected samples at indicated time-points post-infection. Data is representative of three biological replicates, normalized to *MON1* gene expressions, and represented as fold-change relative to WT. Error bars indicates SD. Statistical significance was calculated by Student’s t-test and indicated by asterisks with significance *=*p*<0.05; **=*p*<0.01; ***=*p*<0.001, ns= not significant.

### System development, enrichment and identification of SUMO1-substrates related to defense responses

For robust enrichment and identification of SUMO1/2-substrates modulated during immunity, we utilized two parallel systems for independent validations. First, we generated *srfr1-4::His-SUM1* plants by genetic crosses. These plants express the previously described *His-H89R-SUMO1* (Miller et al. 2010; referred hereafter as *His-SUM1*) transgene in *srfr1-4 sum1-1* background (referred hereafter as *srfr1-4::His-SUM1*) and represents the auto-immune system. Overall, the *srfr1-4::His-SUM1plants* mirror *srfr1-4* pattern in expression levels of *PR1, PR2*, SA-biosynthesis gene *ISOCHORISMATE SYNTHASE 1* (*ICS1*), SUMOylation machineries *SUM2, SAE1a, SAE2, SCE1* and others (Figure S2). A modest increase in *SUM1* expression matching the levels in the parental *His-SUM1* is detected in *srfr1-4::His-SUM1plants*. Because elevated *SUM1* expression is linked to increased SA-responses (van den Burg et al. 2010), the upregulated levels of *SUM3* transcripts likely reflect this phenomenon in the *His-SUM1* transgene containing lines. As observed for *srfr1-4*, SUMO1/2 conjugates accumulate more in *srfr1-4::His-SUM1* extracts. The auto-immune *srfr1-4::His-SUM1* and the *His-SUM1* line represents first system pair for SUMO-enrichments. For the second system, we utilized tissues from mock- (*His-SUM1 Mock*) or *PstDC3000*-challenged (*His-SUM1 Pst*) for 24-hrs for SUMO1/2-enrichments. Tissues from two biological replicates of each system-sets were processed in presence of strong denaturants, subjected to Ni^2+^-NTA affinity chromatography and bound proteins eluted. A schematic flow-chart for processing, enrichment, sample preparation and analysis is shown (Figure 4A). Enrichment efficiencies of SUMO1-conjugates in the eluates were validated by immunoblot with anti-SUMO1 antibodies (Figure 4B). Eluates were then in-gel digested with trypsin and subjected to LC-MS/MS analysis.

**Figure 4.**
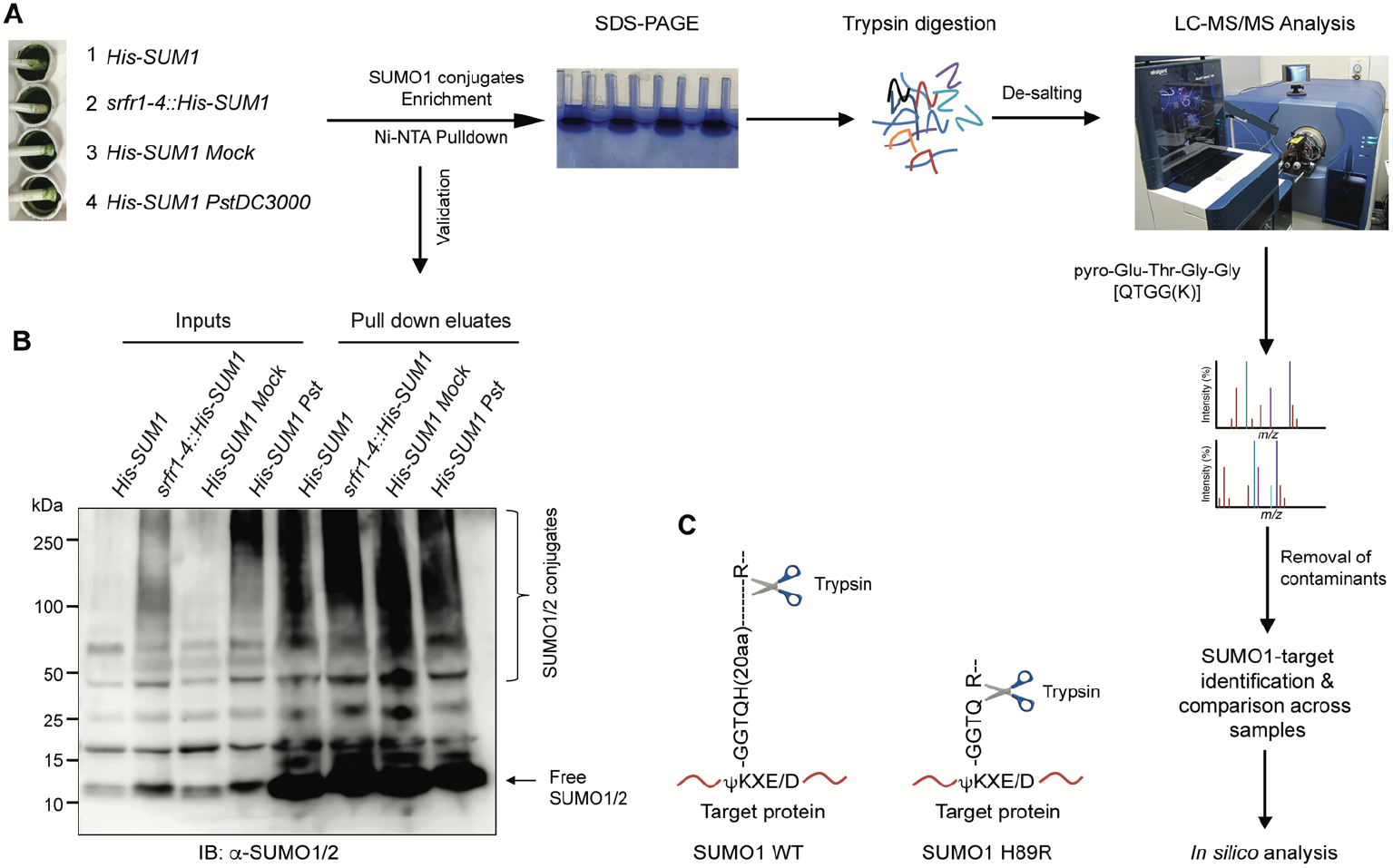
Schematic work-flow of sample preparation and enrichments of SUMO1-conjugates for LC-MS/MS analysis. **(A)** Extracts from indicated plants were enriched for His-SUMO1 via Ni^2+^-NTA pull-down, resolved on SDS-PAGE, in-gel trypsin digested, desalted, LC-MS/MS analysis followed by identification of target proteins and their *in silico* analysis performed as shown stepwise. **(B)** Enrichment of SUMO1-conjugates in the Ni^2+^-NTA eluates detected by immunoblotting with anti-SUMO1/2 antibodies. Positions of SUMO1-conjugates and free-SUMO1 is shown. Migration position of molecular weight standards (in kDa) are indicated. **(C)** Diagrammatic representation of trypsin cleavage sites and respective SUMO1-footprints for wild-type and His-H89R SUMO1 variant (Miller et al. 2010).

The peptide sequences identified were searched against the Arabidopsis proteome database (www.NCBI.nlm.NIH.gov/RefSeq/). A total of 61 background proteins categorized as contaminants from previous reports (Miller et al. 2013; Miller et al. 2010) are detected in our datasets (Table S1). After subtracting these contaminants, a total of 261 SUMO1-SUMOylated proteins including SUMO1, are identified across all samples (Table S2). The Venn diagram distributions of these in the corresponding system-sets and overlaps or distinctions with regard to defense responses, respectively are shown (Figure 5A,B). Of these, 39 and 8 proteins have been previously listed as *bona fide* SUMO1-candidates in heat-stressed *Arabidopsis* (Miller et al. 2013; Miller et al. 2010) or corresponding homologs identified as SUMO-substrates in potato (Colignon et al. 2017a), respectively (Table S2). GPS-SUMO (http://sumosp.biocuckoo.org/online.php) software predicted the presence of at least one or more SUMOylation sites in the primary amino acid sequences of 221 proteins we identified (Table S3). Upon comparison with similar studies, our data reveal greater frequency in enrichment of proteins with predicted SUMOylation sites (Table S4). We identified 8 SUMO1-targets with QTGG footprints on their modified Lysine (Table 1). The MS/MS spectra of these proteins are shown (Figure S5).

**Table 1.**
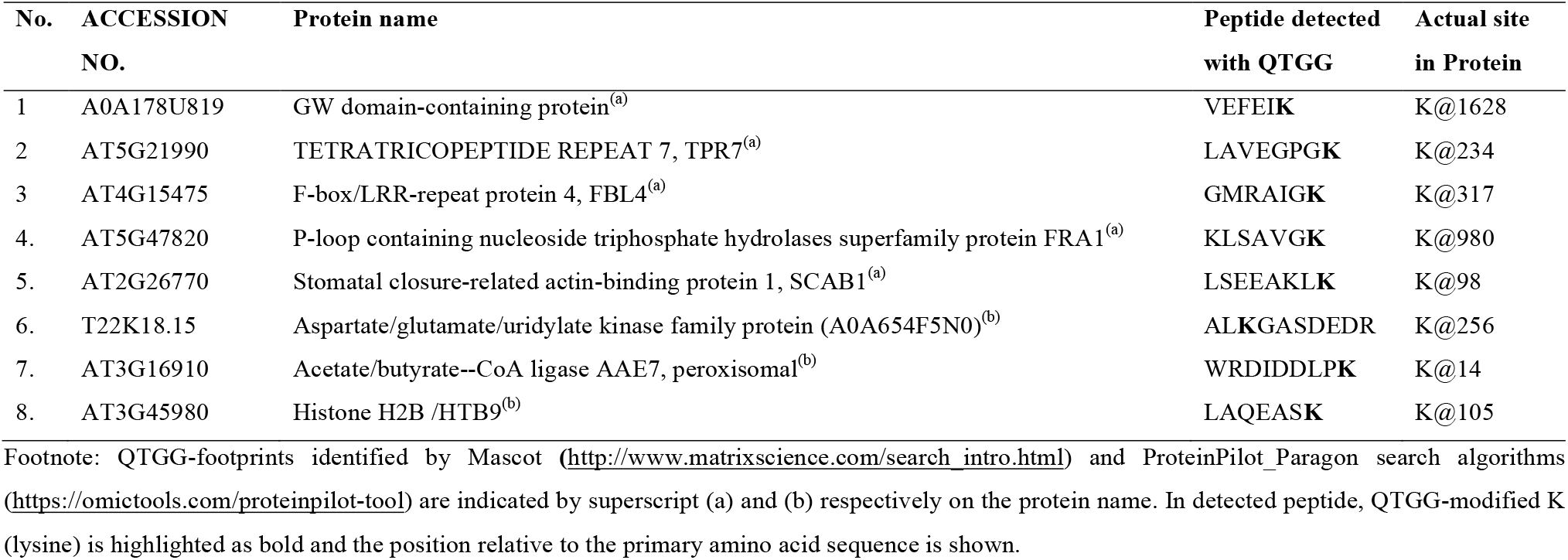
Proteins with QTGG-footprint identified in this study.

**Figure 5.**
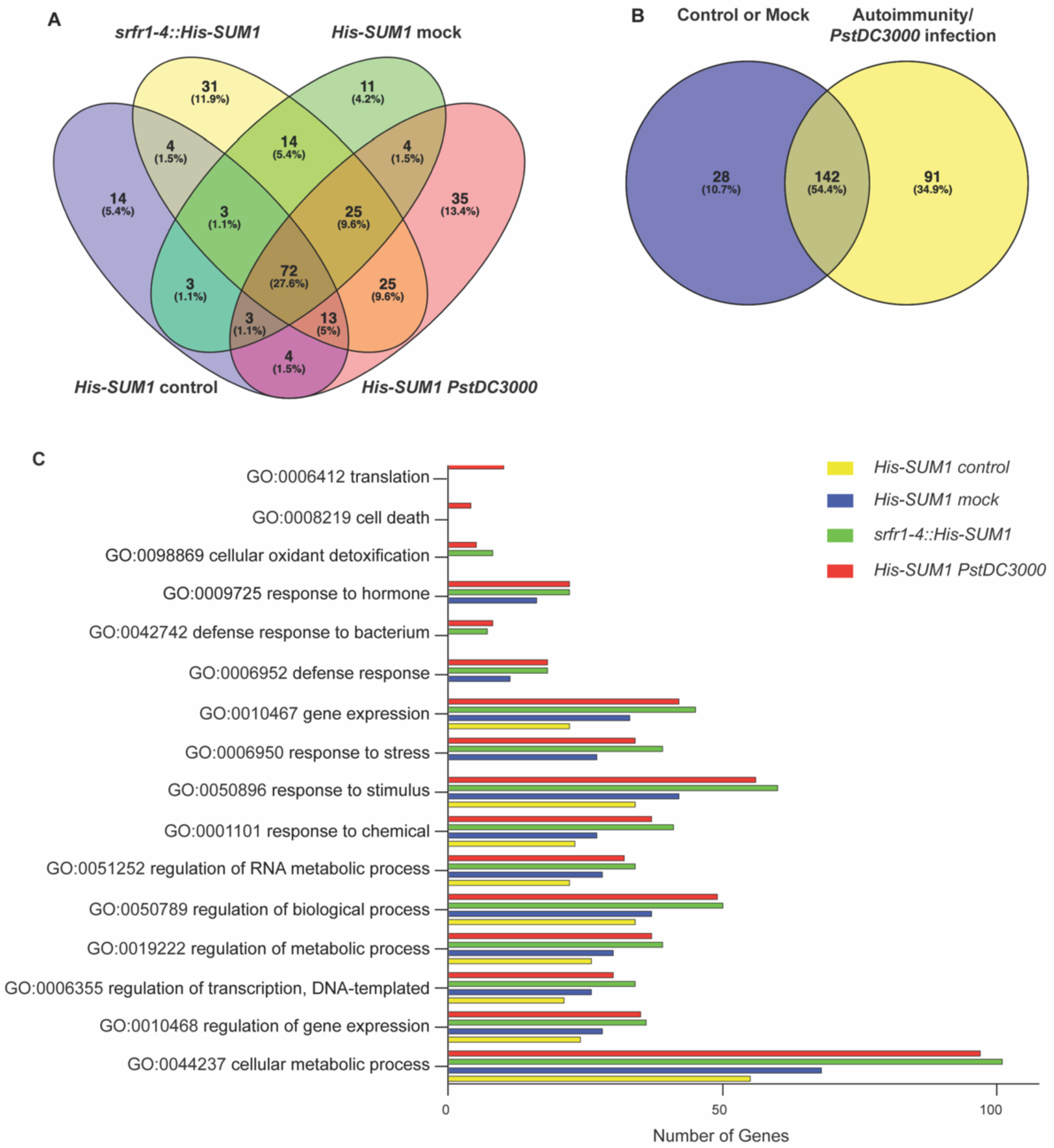
SUMO1-candidates enriched from *srfr1-4* or *PstDC3000* infections categorize predominantly to defense and stress-related processes. **(A)** Venn diagram of enriched SUMO1-candidate numbers and overlaps across different samples. **(B)** Venn diagram of common and distinct SUMO1-substrates identified from *His-SUM1* control or Mock versus *srfr1-4::His-SUM1* or *PstDC3000-infected* samples. **(C)** Representation of selective GO terms with most notable quantitative changes among the different SUMO1-enrichment systems.

### GO analysis and protein-protein interaction (PPI) networks of identified SUMO1-substrates

SUMO1-substrates in our list span across all five arabidopsis chromosomes and also includes few proteins that are chloroplast-encoded (ShinyGO v0.61: Gene ontology enrichment analysis; http://bioinformatics.dstate.edu/go/; Figure S3). Many of the identified proteins have nuclear-assigned functions reinforcing the role of SUMOylation in regulating these processes. Transcription factors populating our list encompass bHLH, AT-Hook, bZIP, WRKY, GATA, and TCPs families suggesting their role in modulating diverse gene expressions. Gene ontology (GO) analysis categorized the identified substrates in 187 GO terms including cellular/metabolic processes, regulation of transcription/translation, RNA-metabolism, general/anti-bacterial defenses, response to various stresses, cellular detoxification, cell death, response to hormone among others (Table S5). Comparisons across the system-sets especially show significant increases in SUMOylation of proteins in the defense-induced samples (*srfr1-4::His-SUM1* and *His-SUM1 PstDC3000*) in comparison to their respective controls (Figure 5C). These also include proteins involved in regular housekeeping processes and implicate their functional adjustments in response to defences. A comprehensive protein-protein interaction (PPI) network of identified proteins was also generated using the STRING database (high confidence of 0.700 with disconnected nodes hidden) (Figure S4). Especially evident in this network is the interconnectivity of the identified SUMO1-substrates with each other. Overall, with the identification of SUMO1-candidates we highlight likely mediators of cellular crosstalks orchestrated by SUMOylation changes during innate immunity.

## DISCUSSION

Several studies have documented the importance of SUMO-modified proteins for basic growth, development and stress adaptations in eukaryotes (Castaño-miquel and Lois, 2016; de Vega et al. 2018; Geiss-Friedlander and Melchior, 2007; Gill, 2004; Srivastava et al. 2016; Verma et al. 2018). In plant systems, barring few reports these remain largely understudied and the identity of differentially SUMOylated candidates connected to immune responses are very few (Colignon et al. 2017a, 2017b; Mazur et al. 2017; Niu et al. 2019; Saleh et al. 2015). Hence it is essential to obtain comprehensive coverage of SUMOylation dynamics and the substrates modulated therein during defenses. We approached these in two independent systems. Evaluation from *PstDC3000*-challenged or auto-immune *srfr1-4* plants allowed identification of SUMO-candidates undergoing both transitory and/or stable modifications during defense responses. Over-representation of proteins related to chromatin architectures such as HDA19/HD1 (HISTONE DEACETYLASE19), Histone H2B /HTB9, HON4, MAM1/HAG4 (HISTONE ACETYLTRANSFERASE of the MYST family 1), WD40, SET, SNF2 among others implicate SUMOylome dynamics in conditioning stimuli-associated rapid transcriptional re-programming. Gain of SUMO-modifications in transcription factors of bHLH, bZIP, AT-hook, TCPs, WRKYs, Trihelix/GATA or ethylene responsive families we especially note, possibly integrate these processes.

Enhancement in global SUMOylome upon application of SA has been reported earlier (Bailey et al. 2015). We demonstrate first that in response to *PstDC3000* infections, progressive increments in SUMO1/2-conjugates occur. Upregulations of *SUM3*, but not *SUM1/2*, transcripts during *PstDC3000* infections or in *srfr1-4* plants are in accordance to enhanced SA-signaling networks suggested earlier since abolishing these via the loss of *EDS1* restores SUMO1/2-conjugate to Col-0 levels (van den Burg et al. 2010). Spatio-temporal adjustments of a SUMOylome is orchestrated by SIZ1 and ESD4 activities (reviewed in Elrouby, 2014). In these modulations, down-regulation of SUMO proteases including *ESD4* during post-stress are especially noted in mammalian systems (Pinto et al. 2012). This is accompanied by enhanced activities of SIZ1 to cause net increase in SUMO-conjugates (Miller et al. 2013). Remarkably, in our transcriptomic data we detect a strong parallel with these observations. Across both our tested systems, progressive increase (upon *PstDC3000* challenge) or enhancement in basal levels (in *srfr1-4*) of transcripts promoting SUMOylation (*SAE2*, *SCE1, SIZ1*) and down-regulation of SUMO-proteases (*ESD4, ELS1*) lend strong support. Perhaps equally significant in our data is that although *SUM1 per se* is not transcriptionally upregulated, existing pools of its mRNAs are more efficiently assembled onto ribosomes for translation during immunity, thus providing necessary protein support to meet SUMO1-conjugation requirements. To our understanding, this is the first demonstration of preferential loading of *SUM1* transcripts onto polyribosomes for translation during autoimmunity. To this end, these results mark rapid deployment of defenses aided through selective translational reprogramming of existing mRNA pools. A significant presence of SUMOylated proteins related to RNA-processes such as RNA binding, end processing and polyadenylation likely drive these mechanisms (Meier, 2012).

Comparative changes in SUMOylome across various stress exposures reveal that instead of newer substrates undergoing covalent modifications, SUMOylation levels on prior-SUMOylated proteins pools are more altered (Miller et al. 2013). In the *His-SUM1* line, 357 SUMO1-targets with abundance of 172 proteins using iTRAQ labelling were earlier reported (Miller et al. 2013, 2010). Our data adds that SUMOylation of several newer proteins nevertheless is considerably promoted during anti-bacterial defenses. We speculate that plants modify specific subset of SUMO-targets appropriate to the nature of stress/stimulus. Of the ~40 heat shock-responsive transcription factors (HSFs) or transcriptional regulators/co-regulators SUMOylated during heat-shock (Miller et al. 2010), only HEAT SHOCK PROTEIN 20 (HSP20) is detected in the *PstDC3000*-infected samples. Nonetheless, several SUMO-substrates we identify are common with previous studies (Table S2) and likely represent the predominant pool of proteins modulated by SUMO. In a comparable degree to Miller et al. (2010), ~85% of the SUMO-substrates we identify have predicted SUMOylation sites thus providing good confidence and coverage in our data (Table S3). Likewise to Miller et al. (2010), QTGG-footprints are detected in the MS-spectra for ~3% (8 out of 261) of the proteins enriched in at least one of the immune-elicited samples.

Although genetically implicated as negative immune regulators, our data clearly support the increasingly evident functional complexities of SUMO1/2 that contrast their classifications (Morrell and Sadanandom, 2019). Of the 29 SUMO-substrates categorized to defense signalling (GO: 0006952), a good proportion includes proteins (such as βCA, RCA, CSD1, MSD1, EULS3, PER34, PUB26, YY1, HSR4, WRKY18, GSTF2) that feature in immunity and are induced in response to pathogens (Table 2; UNIPROT: www.uniprot.org). Among these, UBP1-ASSOCIATED PROTEINS 2A/B (UBPs), CARBONIC ANHYDRASES (βCA, RCA), SUPEROXIDE DISMUTASES (CSD1, MSD1), SERINE HYDROYMETHYL TRANSFERASE (SHMT1), HYPERSENITIVITY-RELATED 4 (HSR4), and WRKY18 have documented roles in regulating HR and ROS responses (Jones et al. 2006; Kim et al. 2008; Moreno et al. 2005; Navarro et al. 2004). Whether upon acquiring SUMO-tags, their activities drive immunity or are supressed to fine-tune balances on energy costs to the host necessitates further mutational studies. Not the least, our data confirms earlier reports that metabolic processes that extend beyond the nuclear locale and include targets in organelles like chloroplast (βCA2, RCA, PETC), endoplasmic reticulum (BG1), mitochondria (SHMT1, MSD1), vacuole (PER34), peroxisomes (PEN2), and extracellular milieu (CYSB) are affected by SUMOs (Colignon et al. 2017a, 2017b; Miller et al. 2010).

**Table 2.**
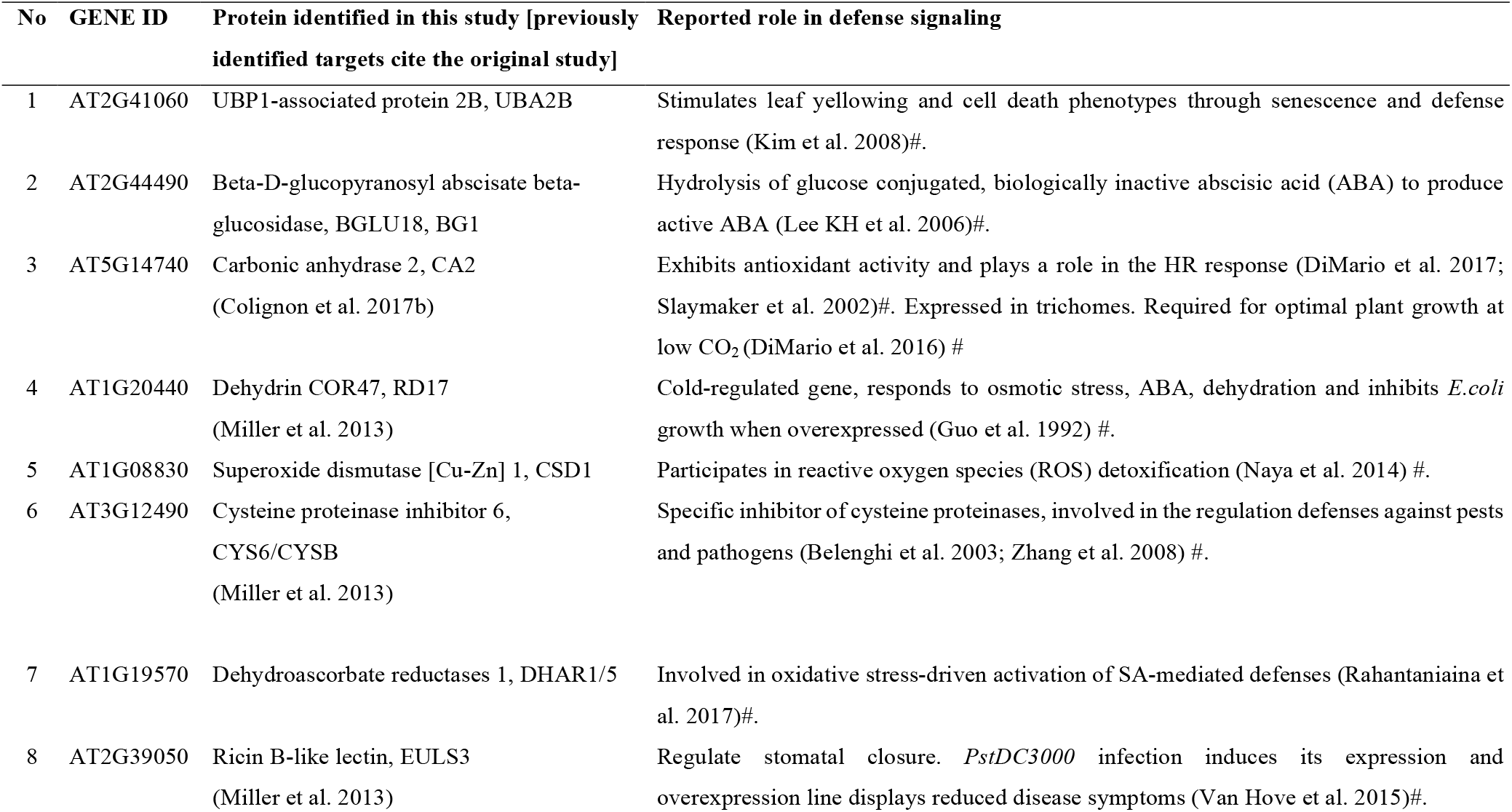
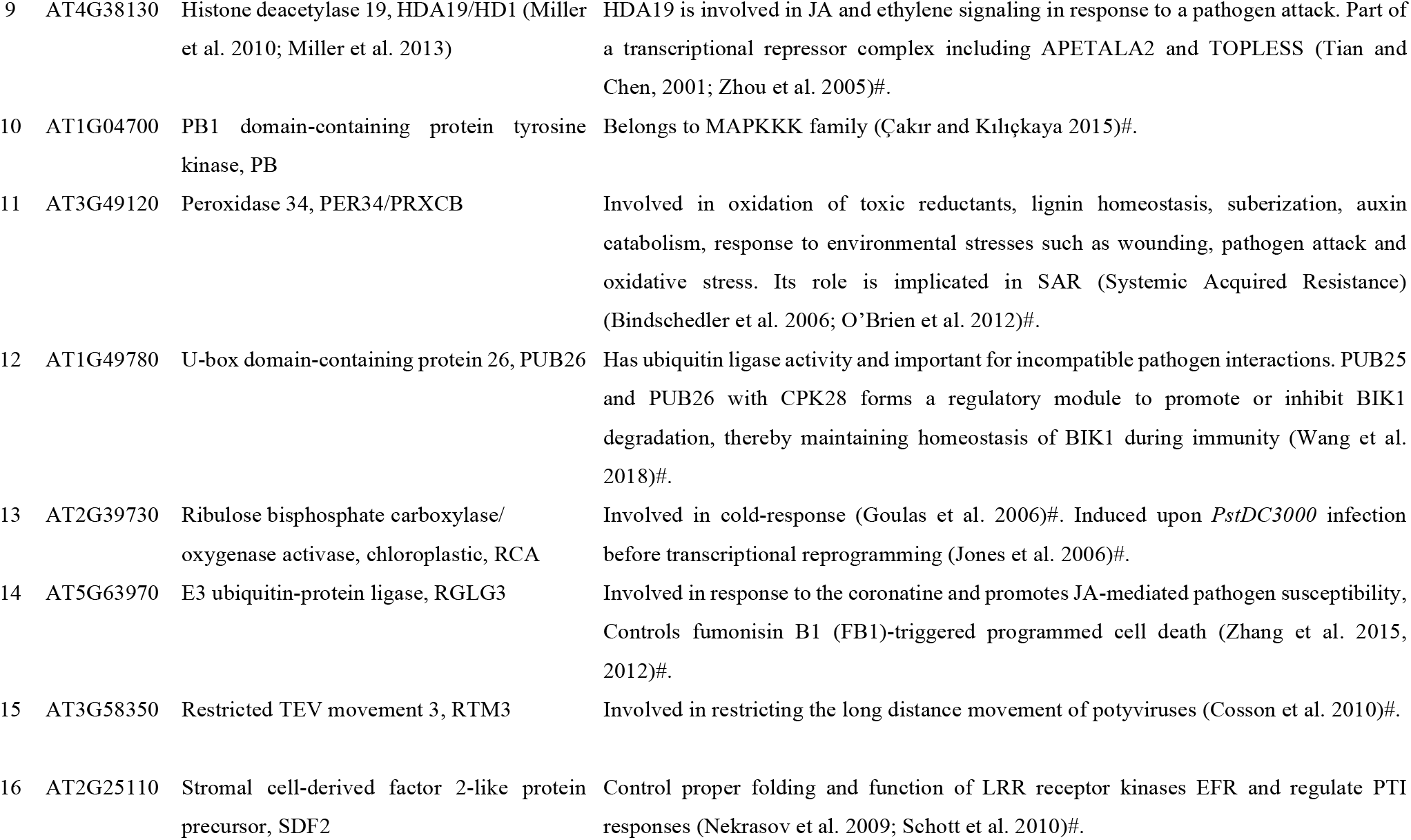
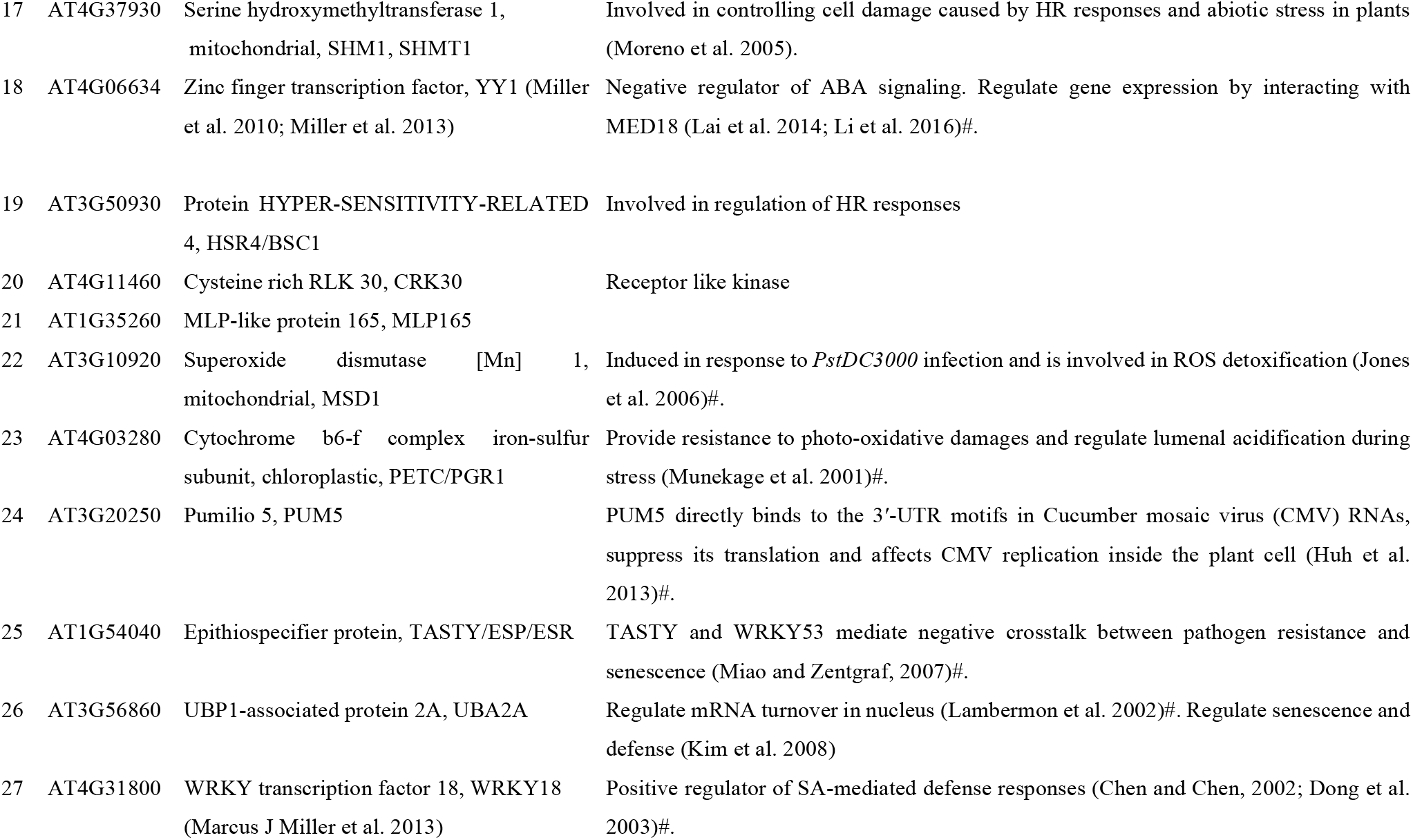
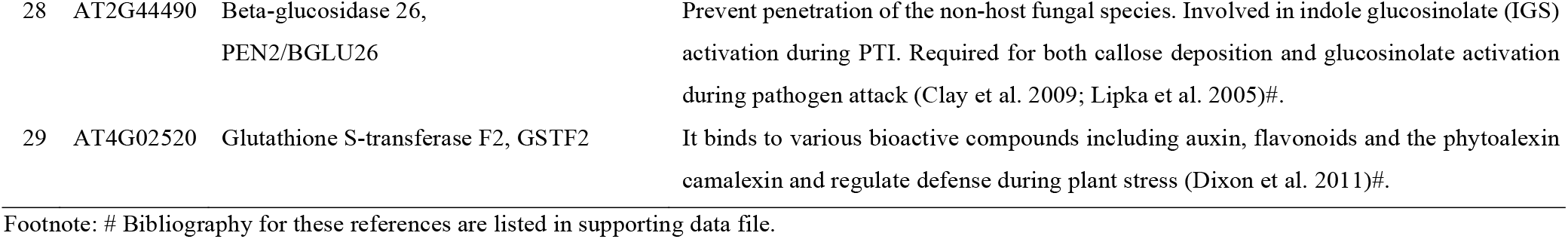
Identified SUMO1-candidates and their reported role in defense signaling (GO:0006952)

| No | GENE ID | Protein identified in this study [previously identified targets cite the original study] | Reported role in defense signaling |
| --- | --- | --- | --- |
| 1 | AT2G41060 | UBP1-associated protein 2B, UBA2B | Stimulates leaf yellowing and cell death phenotypes through senescence and defense response (Kim et al. 2008)#. |
| 2 | AT2G44490 | Beta-D-glucopyranosyl abscisate beta-glucosidase, BGLU18, BG1 | Hydrolysis of glucose conjugated, biologically inactive abscisic acid (ABA) to produce active ABA (Lee KH et al. 2006)#. |
| 3 | AT5G14740 | Carbonic anhydrase 2, CA2<br>(Colignon et al. 2017b) | Exhibits antioxidant activity and plays a role in the HR response (DiMario et al. 2017; Slaymaker et al. 2002)#. Expressed in trichomes. Required for optimal plant growth at low CO<sub>2</sub> (DiMario et al. 2016) # |
| 4 | AT1G20440 | Dehydrin COR47, RD17<br>(Miller et al. 2013) | Cold-regulated gene, responds to osmotic stress, ABA, dehydration and inhibits *E.coli* growth when overexpressed (Guo et al. 1992) #. |
| 5 | AT1G08830 | Superoxide dismutase [Cu-Zn] 1, CSD1 | Participates in reactive oxygen species (ROS) detoxification (Naya et al. 2014) #. |
| 6 | AT3G12490 | Cysteine proteinase inhibitor 6,<br>CYS6/CYSB<br>(Miller et al. 2013) | Specific inhibitor of cysteine proteinases, involved in the regulation defenses against pests and pathogens (Belenghi et al. 2003; Zhang et al. 2008) #. |
| 7 | AT1G19570 | Dehydroascorbate reductases 1, DHAR1/5 | Involved in oxidative stress-driven activation of SA-mediated defenses (Rahantaniaina et al. 2017)#. |
| 8 | AT2G39050 | Ricin B-like lectin, EULS3<br>(Miller et al. 2013) | Regulate stomatal closure. *PstDC3000* infection induces its expression and overexpression line displays reduced disease symptoms (Van Hove et al. 2015)#. |
| 9 | AT4G38130 | Histone deacetylase 19, HDA19/HD1 (Miller et al. 2010; Miller et al. 2013) | HDA19 is involved in JA and ethylene signaling in response to a pathogen attack. Part of a transcriptional repressor complex including APETALA2 and TOPLESS (Tian and Chen, 2001; Zhou et al. 2005)#. |
| 10 | AT1G04700 | PB1 domain-containing protein tyrosine kinase, PB | Belongs to MAPKKK family (Çakır and Kılıçkaya 2015)#. |
| 11 | AT3G49120 | Peroxidase 34, PER34/PRXCB | Involved in oxidation of toxic reductants, lignin homeostasis, suberization, auxin catabolism, response to environmental stresses such as wounding, pathogen attack and oxidative stress. Its role is implicated in SAR (Systemic Acquired Resistance) (Bindschedler et al. 2006; O'Brien et al. 2012)#. |
| 12 | AT1G49780 | U-box domain-containing protein 26, PUB26 | Has ubiquitin ligase activity and important for incompatible pathogen interactions. PUB25 and PUB26 with CPK28 forms a regulatory module to promote or inhibit BIK1 degradation, thereby maintaining homeostasis of BIK1 during immunity (Wang et al. 2018)#. |
| 13 | AT2G39730 | Ribulose biphosphate carboxylase/oxygenase activase, chloroplastic, RCA | Involved in cold-response (Goulas et al. 2006)#. Induced upon *PstDC3000* infection before transcriptional reprogramming (Jones et al. 2006)#. |
| 14 | AT5G63970 | E3 ubiquitin-protein ligase, RGLG3 | Involved in response to the coronatine and promotes JA-mediated pathogen susceptibility, Controls fumonisin B1 (FB1)-triggered programmed cell death (Zhang et al. 2015, 2012)#. |
| 15 | AT3G58350 | Restricted TEV movement 3, RTM3 | Involved in restricting the long distance movement of potyviruses (Cosson et al. 2010)#. |
| 16 | AT2G25110 | Stromal cell-derived factor 2-like protein precursor, SDF2 | Control proper folding and function of LRR receptor kinases EFR and regulate PTI responses (Nekrasov et al. 2009; Schott et al. 2010)#. |
| 17 | AT4G37930 | Serine hydroxymethyltransferase 1, mitochondrial, SHM1, SHMT1 | Involved in controlling cell damage caused by HR responses and abiotic stress in plants (Moreno et al. 2005). |
| 18 | AT4G06634 | Zinc finger transcription factor, YY1 (Miller et al. 2010; Miller et al. 2013) | Negative regulator of ABA signaling. Regulate gene expression by interacting with MED18 (Lai et al. 2014; Li et al. 2016)#. |
| 19 | AT3G50930 | Protein HYPER-SENSITIVITY-RELATED 4, HSR4/BSC1 | Involved in regulation of HR responses |
| 20 | AT4G11460 | Cysteine rich RLK 30, CRK30 | Receptor like kinase |
| 21 | AT1G35260 | MLP-like protein 165, MLP165 |  |
| 22 | AT3G10920 | Superoxide dismutase [Mn] 1, mitochondrial, MSD1 | Induced in response to *PstDC3000* infection and is involved in ROS detoxification (Jones et al. 2006)#. |
| 23 | AT4G03280 | Cytochrome b6-f complex iron-sulfur subunit, chloroplastic, PETC/PGR1 | Provide resistance to photo-oxidative damages and regulate lumenal acidification during stress (Munekage et al. 2001)#. |
| 24 | AT3G20250 | Pumilio 5, PUM5 | PUM5 directly binds to the 3'-UTR motifs in Cucumber mosaic virus (CMV) RNAs, suppress its translation and affects CMV replication inside the plant cell (Huh et al. 2013)#. |
| 25 | AT1G54040 | Epithiospecifier protein, TASTY/ESP/ESR | TASTY and WRKY53 mediate negative crosstalk between pathogen resistance and senescence (Miao and Zentgraf, 2007)#. |
| 26 | AT3G56860 | UBP1-associated protein 2A, UBA2A | Regulate mRNA turnover in nucleus (Lambermon et al. 2002)#. Regulate senescence and defense (Kim et al. 2008) |
| 27 | AT4G31800 | WRKY transcription factor 18, WRKY18 (Marcus J Miller et al. 2013) | Positive regulator of SA-mediated defense responses (Chen and Chen, 2002; Dong et al. 2003)#. |
| 28 | AT2G44490 | Beta-glucosidase 26,<br>PEN2/BGLU26 | Prevent penetration of the non-host fungal species. Involved in indole glucosinolate (IGS) activation during PTI. Required for both callose deposition and glucosinolate activation during pathogen attack (Clay et al. 2009; Lipka et al. 2005)#. |
| 29 | AT4G02520 | Glutathione S-transferase F2, GSTF2 | It binds to various bioactive compounds including auxin, flavonoids and the phytoalexin camalexin and regulate defense during plant stress (Dixon et al. 2011)#. |

763 Footnote: # Bibliography for these references are listed in supporting data file.

In conclusion, we report the dynamic modulation of the Arabidopsis SUMOylome occurring in defense responses. Undoubtedly, phosphorylation, ubiquitination, acetylation and other PTMs that crosstalk with or modulate the efficacy of SUMOylation influence our data (Nukarinen et al. 2017). To further explore these avenues, use of PTM-specific inhibitors and their relative effects on SUMOylome changes are warranted. Substrates destined for SUMOylation-dependent degradation are likely under-represented in our enrichments. SUMOylated CYCLIN T1;5 (CYCT1;5), detected only in control samples, may represent one such target and possibly belong to negative immune regulators. Indeed, *cyct1;5* null mutants are hyper-resistant to Cauliflower mosaic virus (CaMV) infection (Cui et al. 2007). Candidates with QTGG-footprints we identify such as the STOMATAL CLOSURE-RELATED ACTIN BINDING PROTEIN1 (SCAB1) and FRAGILE FIBER 1 (FRA1) regulating stomatal closure or cell wall thickness, respectively have not been documented in defences and are attractive targets to explore further (Zhao et al. 2011; Zhu et al. 2015). Overall, our studies highlight the complexities of signalling networks orchestrated by SUMOylation pathways in plant-pathogen interactions.

## MATERIALS AND METHODS

### Plant materials and growth conditions

The seeds of *Arabidopsis thaliana* mutant lines *srfr1-4* (SAIL_412_E08), *srfr1-4 eds1-2* and *eds1-2*, and transgenic line *sum1-1 sum2-1*::*His-H89R SUM1* were procured from Profs. Walter Gassmann, University of Missouri-Columbia, USA and Richard D. Vierstra, University of Wisconsin, Madison, WI, respectively. All plants used in this study were grown at 22°C with 70% humidity under Short Days conditions (SD; 8 h: 16 h, light: dark) with light intensity of 100 μmol μm^−2^s^−1^. The *srfr1-4::His-H89R SUM1* line was obtained by genetic crossing. The *srfr1-4::His-H89R SUM1* with intact homozygous *sum1-1* T-DNA mutation was identified in F3 population by PCR based genotyping (Primers listed in Table S6).

### *In planta PstDC3000* infection assay

Bacterial infection were performed according to (Kim et al. 2010). Briefly, *PstDC3000* strain at a density of 5×10^6^ cfu/ml^−1^ or mock solution (10mM MgCl_2_) were infiltrated with a needleless syringe into fully expanded rosette leaves of 3-4-weeks-old plants. Tissues were harvested at 0, 3, 6, 12, 24 and 48-hpi (hours post-infection) and were processed separately for total RNA extraction for gene expression studies, for immunoblots with indicated antibodies or for SUMO1-enrichment assays.

### RNA extraction and gene expression analysis by qRT-PCR

Total RNA was isolated from indicated plants with RNAiso Plus (Takara). RNA was reverse transcribed (iScript™ cDNA Synthesis Kit; Bio-Rad) according to manufacturer’s instructions. All qPCR primers used in this study are listed in Table S6. qPCRs were performed in QuantStudio 6 Flex Real-Time PCR system (Applied Biosystems) with 5X HOT FIREPol^®^ EvaGreen^®^ qPCR Mix Plus (ROX) (Solis BioDyne) according to the manufacturer instructions. All qPCR experiments were replicated at least twice with three replicates (n=3). *MON1* (At2g28390) expression was used as internal control (Kim et al. 2010). Relative expression was calculated according to the PCR efficiency^^-ΔΔCt^ formula. Expression differences were normalised to Col-0 and plotted as fold change relative to WT.

### Polysome assay and qRT PCR

Polysome extraction assay was performed according to protocol described by Zanetti et al. (2005) with some minor modifications. Briefly, ~100 mg tissues from three-weeks-old Col-0 and *srfr1-4* plants were ground to a fine powder in liquid nitrogen, resuspended in polysome extraction buffer (PEB) [200mM Tris-HCl, pH 9.0, 25mM EGTA, 200mM KCl, 36mM MgCl_2_, 5mM dithiothreitol (DTT), 50mg/mL cycloheximide, 50mg/mL chloramphenicol, 0.5mg/mL heparin, 1% (v/v) Triton X-100, 1% (v/v) Tween 20, 1% (w/v) Brij-35, 1% (v/v) Igepal CA-630 or NP-40, 2% (v/v) polyoxyethylene, and 1% (w/v) deoxycholic acid] and clarified by centrifugation at 16,000 g for 20 min at 4°C. Supernatant were overlaid on 1.6M sucrose cushion and ultra-centrifuged at 17,0000 g for 18 hr at 4°C. The polysome pellet was resuspended in DEPC (diethylpyrocarbonate)-treated water. RNAs from the pellet fractions were isolated with RNAiso plus, reverse transcribed and qPCR performed as described above.

### Total protein extract preparation and immunoblotting (IB)

For immunoblotting assays, leaf tissues collected at indicated time points/treatments were homogenised in protein extraction buffer [50mM Tris HCl (pH 8.0), 8M Urea, 50mM NaCl, 1% v/v NP-40, 0.5% Sodium deoxycholate, 0.1% SDS and 1mM EDTA] containing 20mM *N*-ethylmaleimide (NEM), 1X plant protease inhibitors cocktail (Sigma Aldrich) and 2% w/v Polyvinyl polypyrrolidone (PVPP). The homogenates were clarified by centrifugation, mixed with 2X Laemmli buffer (0.1M Tris pH 6.8, 20% glycerol, 4% SDS, 100mM DTT and 0.001% Bromophenol blue). Proteins extracts were boiled at 95°C and then separated onto Mini-PROTEAN ^®^ TGX™ precast protein SDS-PAGE (4-15% gradient) gels (BIO-RAD, #4561083). Proteins were transferred onto polyvinylidene fluoride (PVDF) membrane by wet-transfer method. The membrane was blocked with 5% non-fat skim milk and immunoblots were performed with indicated primary antibodies [anti-SUMO1 (Abcam), anti-Actin C3 (Abiocode), anti-PR1 (Agrisera), anti-PR2 (Agrisera)] in 1X TBST at 4°C for overnight. Blots were washed thrice with 1X TBST and then incubated at RT for one hour with appropriate horse-radish peroxidase (HRP)-conjugated secondary antibodies. The blots were then developed using ECL™ Prime western blotting system (GE Healthcare) and visualised in ImageQuant™ LAS 4000 biomolecular imager (GE Healthcare).

### Enrichment and purification of SUMO1-SUMOylated proteins

Purification of SUMOylated proteins were carried out according to Miller et al. (2010) with minor modifications. In brief, 50g plant tissues from 3-weeks-old *His-H89R SUM1* (*His-SUM1*), *srfr1-4::His-SUM1* set, and *His-SUM1* mock- or *PstDC3000*-infected (24-hpi) were grounded and incubated for 1hr at 55°C with 100mL of Extraction Buffer (EB) [100mM Na_2_HPO_4_, 10mM Tris-HCl (pH 8.0), 300mM NaCl, 10mM iodoacetamide (IAA)] containing 7M Guanidine-HCl, 10mM sodium metabisulfite, 2mM PMSF, 10mM NEM and 1X plant protease inhibitor cocktail. The crude extracts were then clarified by centrifugation at 15,000 g. To the supernatant, 10mM imidazole was added and incubated on rotator for overnight at 4°C in the presence of 2.5ml of Ni-NTA beads (Qiagen). The beads were washed with EB containing 6M guanidine-HCl and 0.25% Triton X-100, followed by EB containing 0.25% Triton X-100 and 8M urea. Proteins were eluted with 6M urea, 100mM Na_2_HPO_4_, 10mM Tris-HCl (pH 8.0), 300mM imidazole containing 10mM IAA. The elutes were dialysed (10kDa MWCO) for overnight with two changes of dialysis buffer [100mM Na_2_HPO_4_, 10mM Tris-HCl pH 8.0, 5% Glycerol]. The elutes were concentrated by using Amicon^^®^^ Ultra-4 Centrifugal Filters of 10kDa cut-off according to manufacturer instructions. SUMO1-conjugate enrichments in inputs and eluates were checked by anti-SUMO1 immunoblots. The eluates were then proceeded for trypsin digestion and mass spec analysis.

### Sample preparation for in-gel trypsin digestion

The sample preparation protocol described by Gundry et al. (2010) was followed with minor modifications. The eluates were loaded onto a 6% SDS-PAGE gel and electrophoresed for 15-20 mins for the proteins to enter into the resolving gel (impurities stuck in the wells). The gel was stained with Coomassie brilliant blue (CBB R-250) solution for 30 mins and then de-stained with de-staining solution [50% water, 45% methanol and 5% Glacial acetic acid] for 3-4 hours. Protein bands in the gel were cut into 1mm^3^ pieces using sterile surgical blade and collected in fresh 1.5ml micro-centrifuge tubes. Gel pieces were washed with washing solution [50:50 solution of Acetonitrile (ACN):Water in 50mM Ammonium bicarbonate (ABC)] thrice to remove CBB stain completely. The gel pieces were then dehydrated by adding 500μl of 100% ACN solution for 10 mins, and dried using speed-vac centrifuge at room temperature (RT). Disulfide bonds were reduced by addition of alkylation solution [10mM DTT in 50mM ABC] and incubated at 60°C for 30 mins. Then 100μL of reduction solution [55mM IAA in 50mM ABC] was added and were incubated for 30 mins at RT in dark. Alkylation solution was removed and gel pieces were washed with 500ml washing solution. Gel pieces were dehydrated again with 500μl of 100% ACN for 10 mins. Residual ACN was removed and gel pieces were dried completely using a speed-vac centrifuge.

### In-gel trypsin digestion

Samples were incubated with Trypsin (Promega, USA) at final protease:protein ratio of 1:20 (w/w) for in-gel digestion at 37°C for 16 hrs. Digested peptides were extracted using gradient of 20% to 80% ACN diluted in 0.1% formic acid (FA). Extraction solution having gel pieces were sonicated for 5 mins in ultra-sonicator water bath to achieve maximum recovery of peptides. The pooled extracts were dried using speed-vac centrifuge. The peptides were desalted with Pierce™ C18 Tips (Thermo Scientific), 100μl bed according to manufacturer instructions and peptides were eluted in 100μl of 75% ACN in 0.1% FA. Eluates were dried using speed-vac centrifuge.

### Tandem Mass Spectrometry

Vacuum dried peptides were resuspended in 10μl 2% ACN in 0.1% FA and subjected to MS/MS using TripleTOF^®^ 5600^+^ (ABsciex) mass spectrometer instrument. The mass analyser was attached through the trap and analytical column with specification ChromeXP, 3C18-CL-120, 3 μm, 120 Å and 0.3 × 150 mm respectively. The spray nozzle was connected with electro ionization source to inject the peptide sample in a mass spectrometer. The used flow rate was 5μl/min for the analytical chromatography to separate the peptides in the continuous gradient of elution with the 2-90% acetonitrile for 65 mins total run time. The system uses the solvent composition with mixture of two reservoirs. Reservoir A of 98% water and 2% ACN in 0.1% FA, and B with 98% ACN and 2% water in 0.1% FA was used. The acquisition was executed with conventional data-dependent mode while operating the instrument into an automatic MS and MS/MS fashion. The parent spectra were acquired with the scan range of 350-1250 m/z. The ion source was operated with the following parameters: ISVF = 5500; GS1 = 25; GS2 = 22; CUR = 30. The data dependent acquisition experiments was set to obtain a high resolution TOF– MS scan over a mass range 150-1500 m/z.

### Data processing and analysis

For SUMO1-proteome analysis, raw MS data files were searched and peptide sequences were assigned using the MASCOT software (version 2.7.0, Matrix Science) against the *Arabidopsis thaliana* protein database (www.NCBI.nlm.NIH.gov/RefSeq/). Search parameters included a precursor mass tolerance of 2.5 Da, fragment ion mass tolerance of 0.3 Da and up to 2 missed trypsin cleavages were allowed. The variable modification of Lys (K) residues by SUMO1 [QTGG p343.149184 m/z] was set. Peptide identification was accepted if the threshold was of more than 50% probability (p> 0.5). For further analysis, proteins with minimum one unique peptide with ≥95% confidence level were considered for analysis. Exclusively specific and overlapping proteins in different samples were categorized using Venny 2.1.0 online analysis tool (https://bioinfogp.cnb.csic.es/tools/venny/). For PPI network construction and GO analysis, we used the STRING version 11.0 (https://string-db.org). SUMOylation consensus prediction was performed by GPS-SUMO 2.0 online server (http://sumosp.biocuckoo.org/online.php).

### Statistical Analysis

For all gene-expression experiments, Student’s *t*-test was performed to check significance and denoted by one, two and three asterisks indicating *p-value* <0.05, <0.01, and 0.001, respectively. GraphPad PRISM (version 7.0a) (https://www.graphpad.com/scientific-software/prism/) software was used to perform statistical analysis and for making the graphs. Adobe Illustrator CC (version 2018, https://www.adobe.com/in/products/illustrator.html) software was used to make final images.

## ACKNOWLEDGEMENTS

All authors deeply acknowledge DST-SERB (Grant No: EMR/2016/001899), DBT (Grant No. BT/PR23666/AGIII/103/1039/2018) and UNESCO-Regional Centre for Biotechnology (RCB), Faridabad for providing financial support and central instrumental facilities for this study. We express our gratitude to Prof. Walter Gassmann, University of Missouri, USA, for providing *PstDC3000* strain and seeds of *srfr1-4* and *srfr1-4 eds1-2* mutants used in this study. We are thankful to Prof. Richard Vierstra, University of Wisconsin, Madison, WI, for providing seeds of transgenic line *His-H89R SUM1* used in this study. We also thank Dr. Nirpendra Singh, Scientific Consultant, Advanced Technology Platform Centre (ATPC) and Mr. Nagavara Prasad (Technical assistant, RCB) for their immense help in LC-MS data analysis. KDI acknowledges Department of Biotechnology (DBT), Government of India for providing fellowship during his PhD tenure and KIIT University, Bhubaneswar, India for his PhD registration. SKD thanks RCB for PhD registration and DBT for PhD fellowship. The authors also express gratitude to Dr. Bornali Gohain for her involvement in initial part of this investigations. We also acknowledge the contributions of multiple additional published reports which were not incorporated here due to restriction in citation numbers.

## Author Contributions

SB conceived the research. KDI and SB designed the study. IKD performed all the experiments. SKD helped in *in silico* data analysis. SB performed genetic crosses and supervised the experiments. KDI, and SB analyzed the data and wrote the manuscript.

## SUPPORTING INFORMATION

**Figure S1.** *EDS1* mutation (*eds1-2*) restores enhanced SUMOylome in *srfr1-4* to Col-0 level.

Total protein extracts from indicated plants grown for 3-weeks were immunoblotted with anti-SUMO1/2-antibodies. Migration position of molecular weight standards (in kDa), SUMO1/2-conjugates and free-SUMO1/2 are shown. Actin immunoblot and PonceauS staining of Rubisco demonstrate comparable loading across samples.

**Figure S2.** Validation of *His-SUM1* and *srfr1-4:His-SUM1* systems.

**(A)** Growth phenotypes of indicated plants 4-weeks post-germination.

**(B-C)** SUMOylation enhancements are comparable between *srfr1-4* and *srfr1-4::His-SUM1* (B), or between Col-0 and *His-SUM1* that are mock or *PstDC3000* infected (C).

**(D-F)** Transcriptional changes in *PR1, PR2*, or *ICS1* (D), or SUMOylation-associated genes (E-G) are also comparable between *srfr1-4* and *srfr1-4::His-SUM1*.

**Figure S3.** Chromosome-wise distribution of genes corresponding to the enriched SUMO1-substrate proteins in Arabidopsis genome analysed by ShinyGo (v0.61; http://bioinformatics.sdstate.edu/go/).

**Figure S4.** STRING-based protein-protein interaction (PPI) networks (https://string-db.org) of SUMO1-substrates identified in this study.

**Figure S5.** MS/MS spectra of His-SUMO1 QTGG-footprint on Lys (K) residue of identified peptides.

**Table S1.** List of contaminants/ background proteins not included for analysis in this study.

**Table S2.** List of 261 SUMO1-substrates identified in this study.

**Table S3.** Prediction of SUMOylation sites in identified SUMO1-substrates by GPS-SUMO 2.0 online server (http://sumosp.biocuckoo.org/online.php).

**Table S4.** Predicted frequency of SUMO sites identifications across different studies and in the respective host proteome.

**Table S5.** Gene ontology (GO) enrichments in 261 SUMO1-substrates identified in this study performed by STRING database (https://string-db.org).

**Table S6:** List of primers used in this study.

