## Supplemental Information for "Proteomic analysis of SUMO1-SUMOylome changes during defense elicitation in *Arabidopsis*"

### Supplementary Figures

#### Figure S1

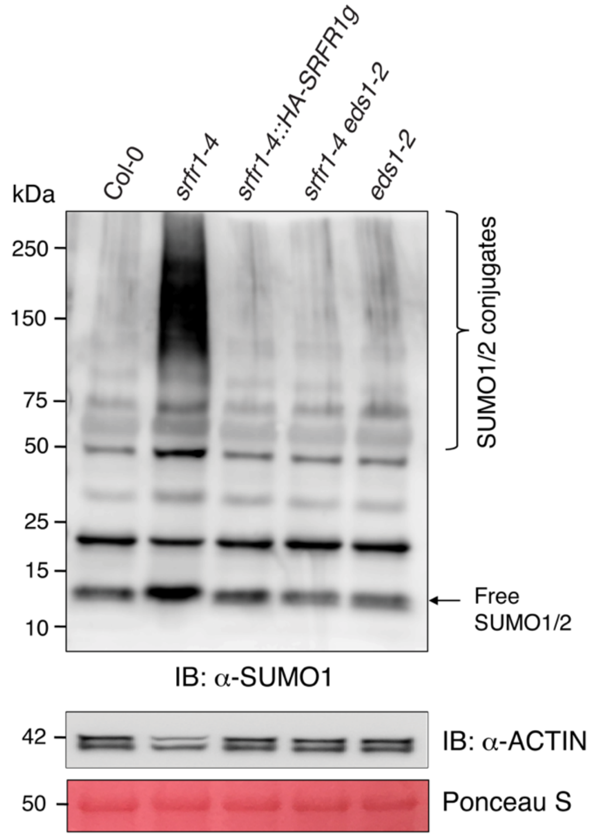


**Figure S1.** ***EDS1*** **mutation (*****eds1-2*****) restores enhanced SUMOylome in** ***srfr1-4*** **to Col-0 level.**

Total protein extracts from indicated plants grown for 3-weeks were immunoblotted with anti-SUMO1/2-antibodies. Migration position of molecular weight standards (in kDa), SUMO1/2-conjugates and free-SUMO1/2 are shown. Actin immunoblot and PonceauS staining of Rubisco demonstrate comparable loading across samples.

### Figure S2

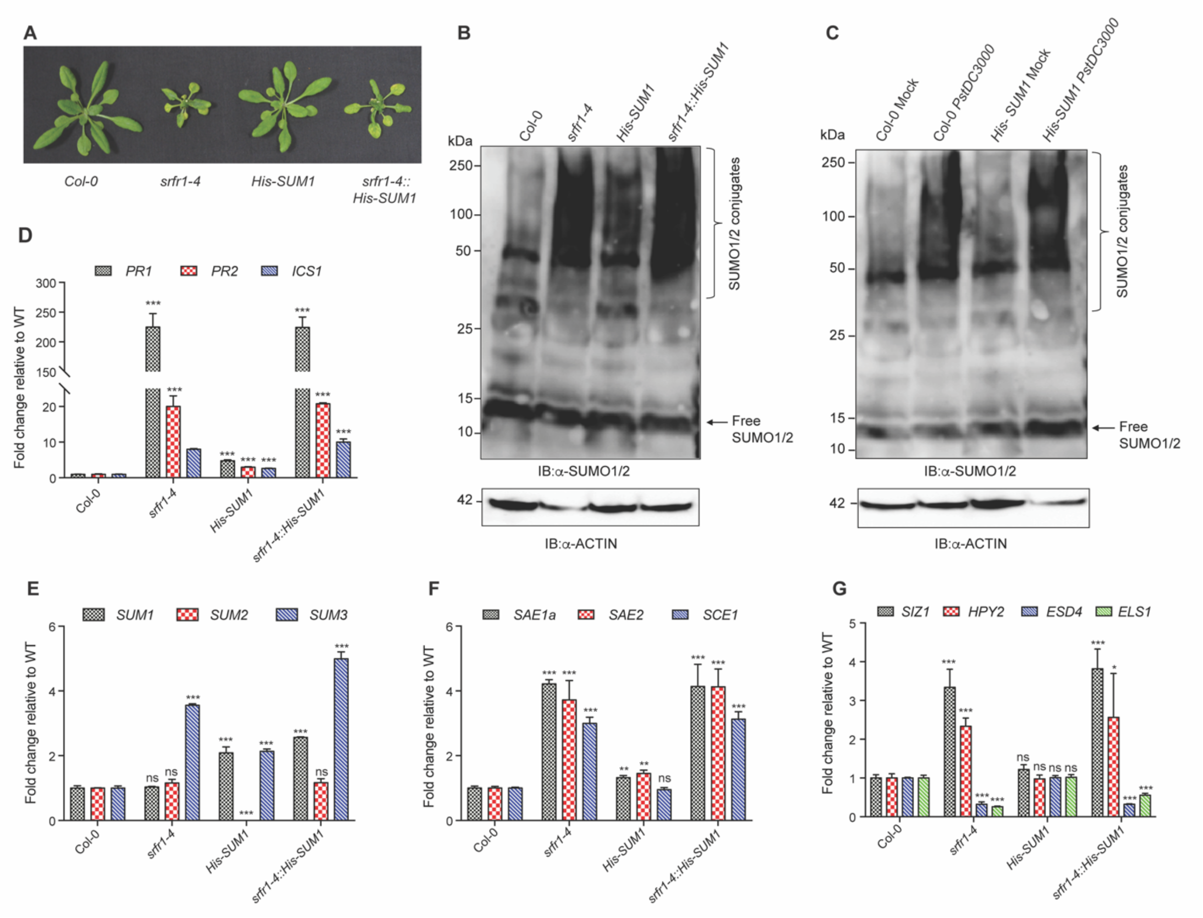


**Figure S2. Validation of *His-SUM1* and *srfr1-4:His-SUM1* systems.**

**(A)** Growth phenotypes of indicated plants 4-weeks post-germination.

**(B-C)** SUMOylation enhancements are comparable between *srfr1-4* and *srfr1-4::His-SUM1* (B), or between Col-0 and *His-SUM1* that are mock or *PstDC3000* infected (C).

**(D-F)** Transcriptional changes in *PR1*, *PR2*, or *ICS1* (D), or SUMOylation-associated genes (E-G) are also comparable between *srfr1-4* and *srfr1-4::His-SUM1*.

**Figure S3**

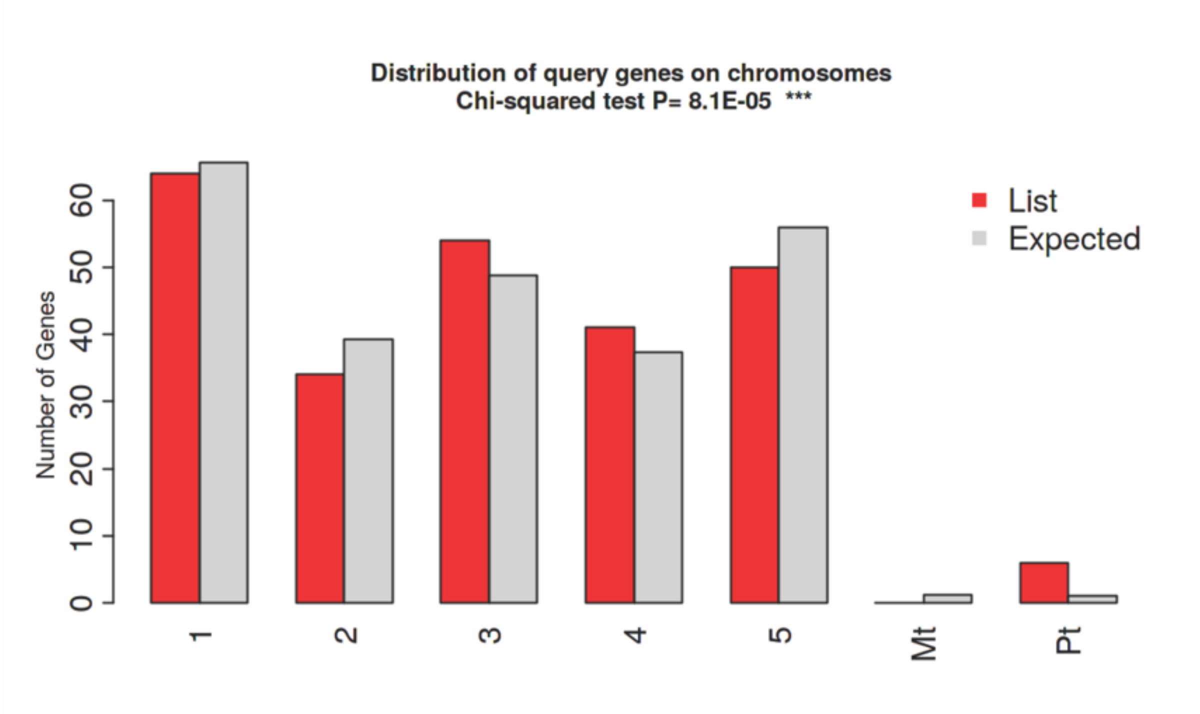


**Figure S3. Chromosome-wise distribution of genes corresponding to the enriched SUMO1-substrate proteins in Arabidopsis genome analysed by ShinyGo (v0.61; http://bioinformatics.sdstate.edu/go/).**

**Figure S4**

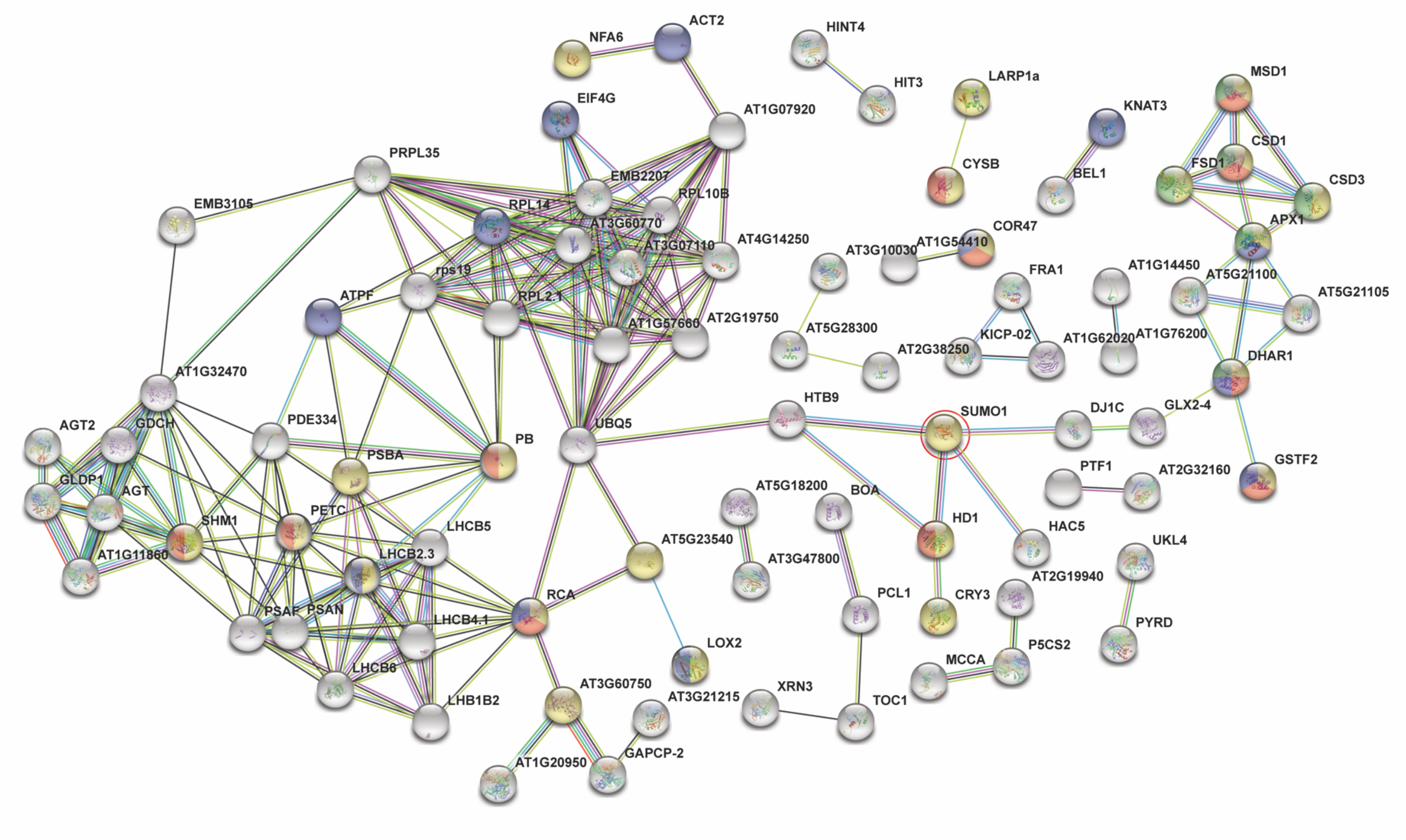


**Figure S4. STRING-based protein-protein interaction (PPI) networks (https://string-db.org) of SUMO1-substrates identified in this study.** (Red-Defense response; Blue-response to hormones, Green-response to reactive oxygen species, Pale yellow-response to stress). Bait protein, SUMO1 is encircled in red.

### Figure S5

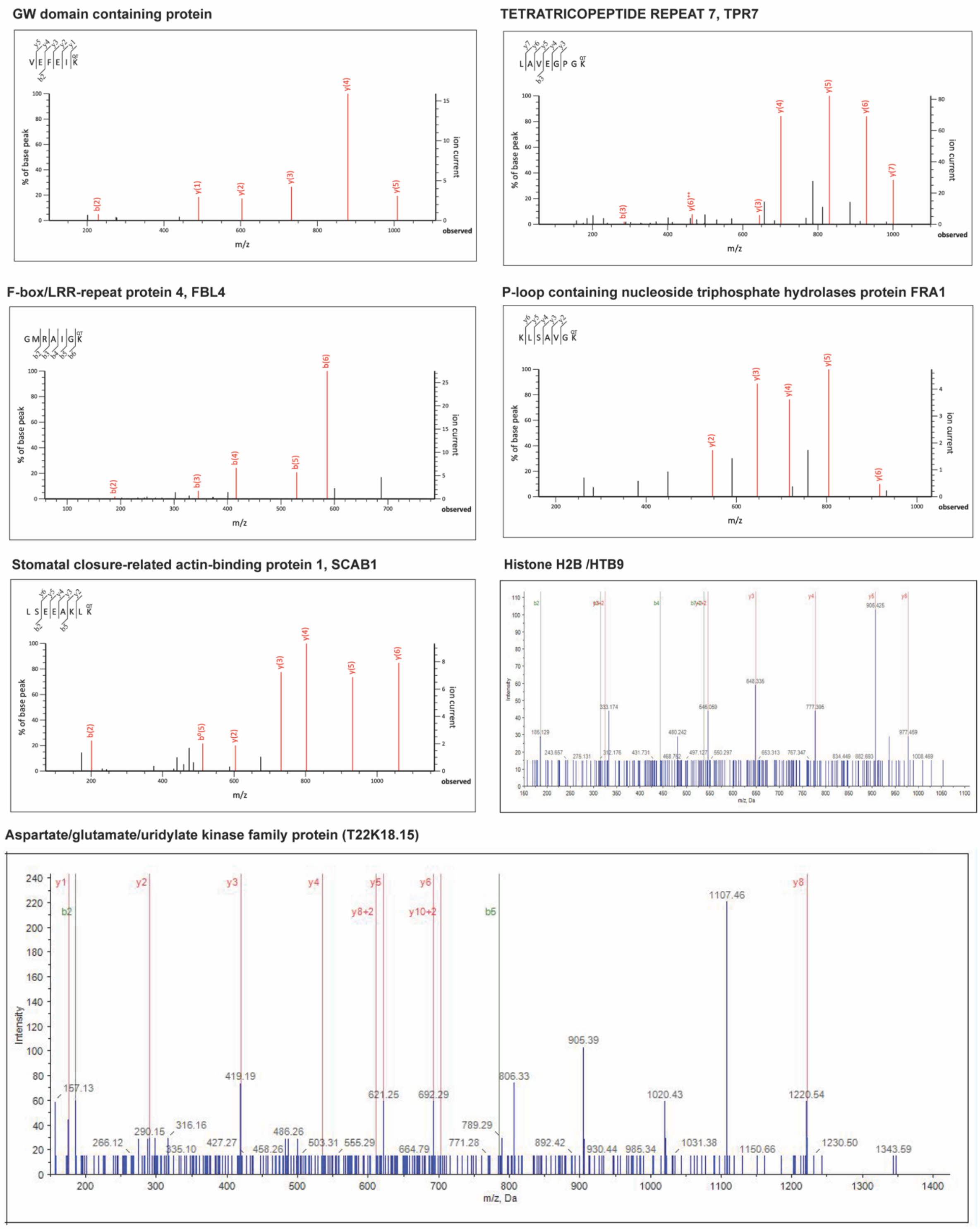


**Figure S5. MS/MS spectra of His-SUMO1 QTGG-footprint on Lys (K) residue of identified peptides.**

### SUPPORTING TABLES

**Table S1. List of contaminants/ background proteins not included for analysis in this study.**

| No | ACCESSION NO | Suggested by | Name | Protein Identified |
|---|---|---|---|---|
| 1 | ATCG00490 | (Miller et al. 2010; 2013) | rbcL | Ribulose bisphosphate carboxylase large chain |
| 2 | AT4G35090 | (Miller et al. 2010; 2013) | CAT2 | Catalase-2 |
| 3 | AT1G20620 | (Miller et al. 2010; 2013) | CAT3 | Catalase-3 |
| 4 | AT1G73840 | (Miller et al. 2010; 2013) | ESP1 | ESP1 |
| 5 | AT3G22380 | (Miller et al. 2010; 2013) | TIC | Protein TIME FOR COFFEE |
| 6 | AT4G28480 | (Miller et al. 2010) | DNAJ | DNAJ heat shock family protein |
| 7 | AT1G70830 | (Miller et al. 2010; 2013) | MLP28 | MLP28 |
| 8 | AT2G34470 | (Miller et al. 2010; 2013) | UREG | Urease accessory protein G |
| 9 | AT1G28290 | (Miller et al. 2010; 2013) | AGP31 | AGP31 |
| 10 | AXX17_AT1G24500 | (Miller et al. 2010) | RPL27AB | RPL27AB |
| 11 | AT4G37830 | (Miller et al. 2010; 2013) | COX6A | Cytochrome c oxidase subunit 6a, mitochondrial |
| 12 | AT5G04710 | (Miller et al. 2010) | | Aspartyl aminopeptidase-like protein |
| 13 | AT2G02070 | (Miller et al. 2010; 2013) | IDD5/IDD11 | Protein indeterminate-domain 5, chloroplastic |
| 14 | AT4G24780 | (Miller et al. 2010; 2013) | Probable pectate lyase 18 | Probable pectate lyase 18 |
| 15 | AT3G09540 | (Miller et al. 2010) | F11F8_12 | Pectate lyase |
| 16 | AT4G09020 | (Miller et al. 2010; 2013) | ISA3 | Isoamylase 3, chloroplastic |
| 17 | AT1G10200 | (Miller et al. 2010; 2013) | WLIM1 | GATA type zinc finger transcription factor family protein |
| 18 | ATCG00580 | (Miller et al. 2010) | psbE | Cytochrome b559 subunit alpha |
| 19 | AT3G15730 | (Miller et al. 2010; 2013) | PLDALPHA1 | Phospholipase D alpha 1 |
| 20 | AT1G03230 | (Miller et al. 2010) | F15K9.16 | Eukaryotic aspartyl protease family protein |

| | | | | |
|---|---|---|---|---|
| 21 | AT5G48900 | (Miller et al. 2010) | PLY20_ARATH | Probable pectate lyase 20 |
| 22 | AT2G18510 | (Miller et al. 2010; 2013) | emb2444 | Putative spliceosome-associated protein |
| 23 | AT2G18020 | (Miller et al. 2010; 2013) | EMB2296 | EMB2296 |
| 24 | AT2G47380 | (Miller et al. 2010) | COX5C1 | Cytochrome c oxidase subunit 5C-1 |
| 25 | AT3G15640 | (Miller et al. 2010) | COX5B-1 | Cytochrome c oxidase subunit 5b-1, mitochondrial |
| 26 | AT1G70890 | (Miller et al. 2010; 2013) | MLP43 | MLP43 |
| 27 | AT5G63180 | (Miller et al. 2010; 2013) | PLY22_ARATH | Probable pectate lyase 22 |
| 28 | AT2G16430 | (Miller et al. 2010; 2013) | PAP10 | Purple acid phosphatase 10 |
| 29 | AT1G04680 | (Miller et al. 2010) | PLY1_ARATH | Probable pectate lyase 1 |
| 30 | AT1G70600 | (Miller et al. 2010) | RPL27AC | 60S ribosomal protein L27a-3 |
| 31 | AT5G66760 | (Miller et al. 2010; 2013) | SDH1-1 | Succinate dehydrogenase [ubiquinone] flavoprotein subunit 1, mitochondrial |
| 32 | AT3G07010 | (Miller et al. 2010; 2013) | PLY8_ARATH | Probable pectate lyase 8 |
| 33 | AT1G70840 | (Miller et al. 2010) | MLP31 | MLP-like protein 31 |
| 34 | AT4G23680 | (Miller et al. 2010; 2013) | AT4g23680/F9D16_150 | Major latex protein-related / MLP-related |
| 35 | AT1G05510 | (Miller et al. 2013) | OBAP1A | Oil body-associated protein 1A |
| 36 | AT1G26630 | (Miller et al. 2010; 2013) | FBR12 | Eukaryotic translation initiation factor 5A |
| 37 | AT1G27430 | (Miller et al. 2013) | GYF domain-containing protein | GYF domain-containing protein At1g27430 |
| 38 | AT1G29350 | (Miller et al. 2013) | DUF1296 | RNA polymerase II degradation factor-like protein (DUF1296) |
| 39 | AT2G28790 | (Miller et al. 2013) | | Pathogenesis-related thaumatin superfamily protein |
| 40 | AT3G04590 | (Miller et al. 2013) | AHL14 | AT-hook motif nuclear-localized protein 14 |
| 41 | AT3G05220 | (Miller et al. 2013) | HIPP34 | Heavy metal transport/detoxification superfamily protein |

| | | | | |
|---|---|---|---|---|
| 42 | AT3G10850 | (Miller et al. 2013) | GLY2 | Hydroxyacylglutathione hydrolase cytoplasmic |
| 43 | AT3G15010 | (Miller et al. 2013) | UBA2C | UBP1-associated protein 2C |
| 44 | AT3G19130 | (Miller et al. 2010; 2013) | RBP47B | Polyadenylate-binding protein RBP47B |
| 45 | AT3G47620 | (Miller et al. 2013) | TCP14 | Transcription factor TCP14 |
| 46 | AT3G55760 | (Miller et al. 2013) | At3g55760 | Uncharacterized protein At3g55760 |
| 47 | AT4G01290 | (Miller et al. 2013) | Chorismate synthase | Chorismate synthase |
| 48 | AT5G08450 | (Miller et al. 2013) | HDC1 | RXT3-like protein HDC1 |
| 49 | AT4G23670 | (Miller et al. 2013) | | Bet_v_1 domain-containing protein |
| 50 | AT4G24680 | (Miller et al. 2013) | MOS1 | Protein MODIFIER OF SNC1 1 |
| 51 | AT4G26750 | (Miller et al. 2013) | LIP5 | Protein HOMOLOG OF MAMMALIAN LYST-INTERACTING PROTEIN 5 |
| 52 | AT4G27320 | (Miller et al. 2013) | PHOS34 | PHOS34 |
| 53 | AT5G12230 | (Miller et al. 2013) | MED19A | Mediator of RNA polymerase II transcription subunit 19a |
| 54 | AT5G13780 | (Miller et al. 2013) | NAA10 | N-terminal acetyltransferase A complex catalytic subunit |
| 55 | AT2G01190 | (Miller et al. 2013) | PDE331 | PIGMENT DEFECTIVE 331 |
| 56 | AT5G54430 | (Miller et al. 2013) | PHOS32 | Universal stress protein PHOS32 |
| 57 | AT3G50370 | (Miller et al. 2013) | | Uncharacterized protein At3g50370 |
| 58 | AT3G17240 | (Miller et al. 2013) | LPD2 | Dihydrolipoyl dehydrogenase 2, mitochondrial |
| 59 | AT1G48030 | (Miller et al. 2010; 2013) | mtLPD1 | Dihydrolipoyl dehydrogenase 1, mitochondrial |
| 60 | AT1G70850 | (Miller et al. 2010; 2013) | MLP34 | MLP-like protein 34 |
| 61 | AT5G39570 | (Miller et al. 2013) | At5g39570 | Uncharacterized protein At5g39570 |

**Table S2. List of 261 SUMO1-substrates identified in this study**

| No | Previously identified as SUMO1 substrate | AT# | Protein Name | *His-SUM1* | *srfr1-4::HisSUM1* | *His-SUM1* Mock | His-SUM1 *PstDC3000* |
|---|---|---|---|---|---|---|---|
| 1 | | AT4G26840 | His-H89R SUMO1, | Y | Y | Y | Y |
| 2 | (Miller et al. 2013) | AT2G39050 | Ricin B-like lectin EULS3 | Y | Y | Y | Y |
| 3 | | AT4G24870 | MSP domain-containing protein | Y | Y | Y | Y |
| 4 | (Miller et al. 2010; 2013) | AT1G74560 | NAP1 related protein 1, NRP1 | Y | Y | Y | Y |
| 5 | (Miller et al. 2013) | AT1G11860 | Aminomethyltransferase | Y | Y | Y | Y |
| 6 | | AT3G17450 | BED-type domain-containing protein | Y | Y | Y | Y |
| 7 | | AT4G39660 | Alanine:glyoxylate aminotransferase 2 | Y | Y | Y | Y |
| 8 | | AT4G14250 | Ribosomal_L2_C domain-containing protein | Y | Y | Y | Y |
| 9 | (Miller et al. 2013) | AT1G62020 | Coatomer subunit alpha COP1 | Y | Y | Y | Y |
| 10 | | AT4G31340 | Myosin heavy chain-like protein | Y | Y | Y | Y |
| 11 | (Miller et al. 2013) | AT4G33010 | Glycine cleavage system P protein, GLDP1 | Y | Y | Y | Y |
| 12 | | AT2G35940 | EDA29/BLH1 | Y | Y | Y | Y |
| 13 | | AT1G19570 | DHAR5/DHAR1 | Y | Y | Y | Y |
| 14 | (Miller et al. 2010; 2013) | AT1G80480 | Plastid transcriptionally active 17, PTAC17 | Y | Y | Y | Y |
| 15 | (Miller et al. 2010; 2013) | AT2G43970 | La-related protein 6B, Transcriptional regulator, LARP6B | Y | Y | Y | Y |
| 16 | | AT1G04700 | PB1 domain-containing protein | Y | Y | Y | Y |

| | | | | | | | |
|---|---|---|---|---|---|---|---|
| 17 | | AT2G25110 | Stromal cell-derived factor 2-like protein precursor, SDF2 | Y | Y | Y | Y |
| 18 | | AT4G02290 | Endoglucanase 17, GH9B13 | Y | Y | Y | Y |
| 19 | (Colignon et al. 2017a) | AT3G60750 | Transketolase-1, TKL-1 | Y | Y | Y | Y |
| 20 | | AT3G28920 | Zinc-finger homeodomain protein 9, ZHD9/HB34 | Y | Y | Y | Y |
| 21 | (Miller et al. 2013) | AT3G12490 | Cysteine proteinase inhibitor 6, CYS6/CYSB | Y | Y | Y | Y |
| 22 | | AT1G52730 | Transducin/WD40 repeat-like superfamily protein, FY | Y | Y | Y | Y |
| 23 | | AT3G20820 | LRRNT_2 domain-containing protein | Y | Y | Y | Y |
| 24 | | AT3G01670 | Protein SIEVE ELEMENT OCCLUSION A, SEOR2/SEOA | Y | Y | Y | Y |
| 25 | (Miller et al. 2013) | AT3G42170 | BED zinc finger and domain-containing protein DAYSLEEPER/HAT | Y | Y | Y | Y |
| 26 | (Miller et al. 2013) | AT1G14440 | Zinc-finger homeodomain protein 4, ZHD4 | Y | Y | Y | Y |
| 27 | | AT3G47810 | Vacuolar protein sorting-associated protein 29, VSP29, MAG1 | Y | Y | Y | Y |
| 28 | | AT5G64460 | Phosphoglycerate mutase-like protein 1 | Y | Y | Y | Y |
| 29 | (Colignon et al. 2017a) | AT3G52720 | alpha carbonic anhydrase (CAH1), chloroplastic, ACA1/C, AH1 | Y | Y | Y | Y |
| 30 | | AT5G13780 | N-terminal acetyltransferase A complex catalytic subunit NAA10 | Y | Y | Y | Y |
| 31 | | AT5G12010 | Uncharacterized protein F14F18_180 | Y | Y | Y | Y |

| | | | | | | |
|---|---|---|---|---|---|---|
| 32 | AT1G31160 | HISTIDINE TRIAD NUCLEOTIDE-BINDING 2, HINT2 | Y | Y | Y | Y |
| 33 | AT1G54410 | Dehydrin HIRD11 | Y | Y | Y | Y |
| 34 | AT5G57610 | Protein kinase domain-containing protein | Y | Y | Y | Y |
| 35 | A0A178U819 | GW domain-containing protein | Y | Y | Y | Y |
| 36 | AT2G47710 | Adenine nucleotide alpha hydrolases-like superfamily protein | Y | Y | Y | Y |
| 37 | AT5G18200 | ADP-glucose phosphorylase At5g18200 | Y | Y | Y | Y |
| 38 | C24_LOCUS3675 | Pectinesterase domain-containing protein C24_LOCUS3675 | Y | Y | Y | Y |
| 39 | AT1G03090 | Methylcrotonoyl-CoA carboxylase subunit alpha, mitochondrial, MCCA | Y | Y | Y | Y |
| 40 | AT5G23300 | Dihydroorotate dehydrogenase (quinone), mitochondrial, PYRD | Y | Y | Y | Y |
| 41 | AT1G13450 | Trihelix transcription factor GT-1 | Y | Y | Y | Y |
| 42 | AT1G15125 | Uncharacterized protein At1g15125 | Y | Y | Y | Y |
| 43 | AT5G24850 | Cryptochrome DASH, chloroplastic/mitochondrial, CRY3 | Y | Y | Y | Y |
| 44 | AT5G61550 | U-box domain-containing protein 52, PUB52 | Y | Y | Y | Y |
| 45 | AT2G01210 | Receptor protein kinase-like protein ZAR1 | Y | Y | Y | Y |
| 46 | AT1G59710 | Actin cross-linking protein (DUF569) | Y | Y | Y | Y |
| 47 | AT3G50240 | Kinesin-like protein KICP-02 | Y | Y | Y | Y |

| | | | | | | | |
|---|---|---|---|---|---|---|---|
| 48 | (Miller et al. 2010) | AT1G76510 | ARID/BRIGHT DNA-binding domain-containing protein | Y | Y | Y | Y |
| 49 | (Colignon et al. 2017a) | AT1G16300 | Glyceraldehyde-3-phosphate dehydrogenase GAPCP2 | Y | Y | Y | Y |
| 50 | (Miller et al. 2010; 2013) | AT4G06634 | Zinc finger transcription factor YY1 | Y | Y | Y | Y |
| 51 | | AT5G21990 | Tetratricopeptide repeat (TPR) containing protein 7, OEP61, TRP7 | Y | Y | Y | Y |
| 52 | | AT3G23070 | CRM-domain containing factor CFM3A | Y | Y | Y | Y |
| 53 | | AT1G45150 | Alpha-1,6-mannosyl-glycoprotein 2-beta-N-acetylglucosaminyltransferase | Y | Y | Y | Y |
| 54 | | AT4G30310 | FGGY family of carbohydrate kinase | Y | Y | Y | Y |
| 55 | | AT2G43160 | ENTH domain-containing protein | Y | Y | Y | Y |
| 56 | | AT3G29200 | Chorismate mutase 1, chloroplastic, CM1 | Y | Y | Y | Y |
| 57 | | AT1G75240 | Zinc-finger homeodomain protein 5, ZHD5, HB33 | Y | Y | Y | Y |
| 58 | (Miller et al. 2013) | AT3G21215 | RNA-binding (RRM/RBD/RNP motifs) family protein | Y | Y | Y | Y |
| 59 | | AT1G06240 | Uncharacterized protein At1g06240 | Y | Y | Y | Y |
| 60 | | AT3G55610 | Delta 1-pyrroline-5-carboxylate synthase 2, P5CS2 | Y | Y | Y | Y |
| 61 | | AT4G13990 | Exostosin domain-containing protein Probable xyloglucan galactosyltransferase, GT14 | Y | Y | Y | Y |
| 62 | | AT1G78260 | RNA-binding (RRM/RBD/RNP motifs) family protein | Y | Y | Y | Y |

| | | | | | | | |
|---|---|---|---|---|---|---|---|
| 63 | | AT3G45140 | Lipoxygenase 2, chloroplastic, LOX2 | Y | Y | Y | Y |
| 64 | | AT4G34020 | Protein DJ-1 homolog C, DJ1C | Y | Y | Y | Y |
| 65 | | AT5G47950 | Acetyl-CoA:benzylalcohol acetyltranferase-like protein | Y | Y | Y | Y |
| 66 | | AT5G12290 | Protein DGS1, mitochondrial | Y | Y | Y | Y |
| 67 | | AT5G45190 | Cyclin-T1-5, CYCT1-5 | Y | N | N | N |
| 68 | | AT1G23010 | Multicopper oxidase LPR1 | Y | N | N | N |
| 69 | | AT5G44120 | 12S seed storage protein CRA1, CRU1 | Y | N | N | N |
| 70 | | C24_LOCUS 3590 | SET domain-containing protein C24_LOCUS3590 | Y | N | N | N |
| 71 | | AT5G58350 | Probable serine/threonine-protein kinase WNK4, ZIK2 | Y | N | N | N |
| 72 | | AT4G12290 | Amine oxidase | Y | N | N | N |
| 73 | | AT5G10250 | BTB/POZ domain-containing protein DOT3, Phototropic-responsive NPH3 family protein | Y | N | N | N |
| 74 | | AT1G20950 | Pyrophosphate--fructose 6-phosphate 1-phosphotransferase subunit alpha, PFP-ALPHA1 | Y | N | N | N |
| 75 | | AT2G45880 | Beta-amylase 7, BAM7 | Y | N | N | N |
| 76 | (Miller et al. 2013) | AT1G20440 | Dehydrin COR47, RD17 | Y | Y | Y | N |
| 77 | | AXX17_AT 5G09370 | Actin-7, ACT7 | Y | N | N | N |

| | | | | | | | |
|---|---|---|---|---|---|---|---|
| 78 | | AT5G47820 | Kinesin-like protein KIN-4A, P-loop containing nucleoside triphosphate hydrolases superfamily protein FRA1, KIN4A | Y | N | N | N |
| 79 | (Miller et al. 2010; 2013) | AT3G06480 | DEAD box RNA helicase family protein, RH40 | Y | N | N | N |
| 80 | | AT5G17430 | AP2-like ethylene-responsive transcription factor BBM | Y | N | N | N |
| 81 | | AT1G54040 | Epithiospecifier protein TASTY | Y | N | N | N |
| 82 | | AT1G35260 | MLP-like protein 165, MLP165 | N | N | Y | N |
| 83 | | AT2G32690 | Glycine-rich protein 23, GRP23 | N | N | Y | N |
| 84 | | AXX17_ATUG01340 | Uncharacterized protein AXX17_ATUG01340 | N | N | Y | N |
| 85 | | AT3G13960 | Growth-regulating factor 5, GRF5 | N | N | Y | N |
| 86 | | AT3G45190 | SIT4 phosphatase-associated family protein | N | N | Y | N |
| 87 | | AT3G07750 | 3'-5'-exoribonuclease family protein MLP3.20 | N | N | Y | N |
| 88 | | AT2G44490 | Beta-glucosidase 26, BGLU26, PEN2 | N | N | Y | N |
| 89 | | AT2G38750 | Annexin 4, ANNAT4 | N | N | Y | N |
| 90 | | AT4G11460 | Cysteine rich RLK (RECEPTOR-like protein kinase) 30, CRK30 | N | N | Y | N |
| 91 | | AT2G40620 | bZIP transcription factor 18, BZIP18 | N | N | Y | N |
| 92 | | AT1G77620 | p-loop containing nucleoside triphosphate hydrolases superfamily protein At1g77620 | N | N | Y | N |
| 93 | | AT1G69310 | WRKY DNA-binding protein 57, WRKY57 | N | Y | N | N |

| | | | | | | | |
|---|---|---|---|---|---|---|---|
| 94 | (Miller et al. 2010) | AT5G41410 | Plant Homeobox (POX) protein BEL1 homolog | N | Y | N | N |
| 95 | (Colignon et al. 2017b) | AT3G27690 | Chlorophyll a-b binding protein, chloroplastic LHCB2.3 | N | Y | N | N |
| 96 | | AT1G76200 | NADH dehydrogenase [ubiquinone] 1 beta subcomplex subunit 2 At1g76200 | N | Y | N | N |
| 97 | | AT2G38250 | Trihelix transcription factor GT-3b | N | Y | N | N |
| 98 | | AT4G38220 | Peptidase M20/M25/M40 family protein, AQI | N | Y | N | N |
| 99 | | AT1G68780 | At1g68780 Uncharacterized protein F14K14.11 | N | Y | N | N |
| 100 | | AT1G13350 | Protein kinase domain-containing protein At1g13350 | N | Y | N | N |
| 101 | | AT5G64040 | Photosystem I reaction center subunit N, chloroplastic, PSAN | N | Y | N | N |
| 102 | | AT2G31790 | UDP-glycosyltransferase 74C1, UGT74C1 | N | Y | N | N |
| 103 | | AT5G14740 | Carbonic anhydrase 2, CA2 | N | Y | N | N |
| 104 | | ATCG00020 | Photosystem II protein D1, psbA | N | Y | N | N |
| 105 | | AT1G08830 | Superoxide dismutase [Cu-Zn] 1, CSD1 | N | Y | N | N |
| 106 | | AT4G25940 | Putative clathrin assembly protein | N | Y | N | N |
| 107 | | AT5G51540 | Mitochondrial intermediate peptidase, mitochondrial OCT1 | N | Y | N | N |
| 108 | | AT1G57660 | 60S ribosomal protein L21-2 RPL21E | N | Y | N | N |
| 109 | | AT4G08780 | Peroxidase 38, PER38 | N | Y | N | N |

| | | | | | | | |
|---|---|---|---|---|---|---|---|
| 110 | | AT1G08290 | WPP domain-interacting protein 3, Zinc finger protein WIP3; Probable transcriptional regulator, WIP3 | N | Y | N | N |
| 111 | | AN1_LOCUS20462 | Uncharacterized protein AN1_LOCUS20462 | N | Y | N | N |
| 112 | | AT3G47800 | Aldose 1-epimerase T23J7.130 | N | Y | N | N |
| 113 | | AT1G49780 | U-box domain-containing protein 26, PUB26 | N | Y | N | N |
| 114 | (Miller et al. 2010) | AT2G26230 | Uricase At2g26230 | N | Y | N | N |
| 115 | | AT5G18100 | Superoxide dismutase [Cu-Zn] 3, CSD3 | N | Y | N | N |
| 116 | | AT4G24060 | Dof zinc finger protein DOF4.6 | N | Y | N | N |
| 117 | (Miller et al. 2010; 2013) | AT4G25100 | Superoxide dismutase, FSD1 | N | Y | N | N |
| 118 | | AT5G09410 | Calmodulin-binding transcription activator 5 CAMTA5/EICBP.B | N | Y | N | N |
| 119 | | ATCG00780 | 50S ribosomal protein L14, chloroplastic rpl14 | N | Y | N | N |
| 120 | | AT4G29390 | 40S ribosomal protein S30 RPS30A | N | Y | N | N |
| 121 | | AT3G49120 | Peroxidase 34 , PER34/RXCB | N | Y | N | N |
| 122 | | AT3G12980 | Histone acetyltransferase HAC5 | N | Y | N | N |
| 123 | | AT4G02470 | AAA-type ATPase family protein At4g02470 | N | Y | N | N |
| 124 | (Miller et al. 2013) | AT1G43170 | 60S ribosomal protein L3-1 ARP1/RPL3A/EMB2207 | N | N | N | Y |
| 125 | (Colignon et al. 2017b) | AT1G58280 | Phosphoglycerate mutase family protein At1g58280, F19C14.10 | N | N | N | Y |

| | | | | | | | |
|---|---|---|---|---|---|---|---|
| 126 | | AT3G07110 | 60S ribosomal protein L13a-1, RPL13AA | N | N | N | Y |
| 127 | | AT2G41940 | Zinc finger protein 8, ZFP8 | N | N | N | Y |
| 128 | | AT3G60770 | 40S ribosomal protein S13-1, RPS13A | N | N | N | Y |
| 129 | | AT5G21160 | LA RNA-binding protein 1A, LARP1a | N | N | N | Y |
| 130 | | AT2G31070 | Transcription factor TCP10 | N | N | N | Y |
| 131 | | AT2G42840 | Protodermal factor 1, PDF1 | N | N | N | Y |
| 132 | | AT1G14830 | Dynamin-related protein 1C, DRP1C | N | N | N | Y |
| 133 | | AT5G56980 | Pathogen-associated molecular patterns-induced protein A70 | N | N | N | Y |
| 134 | | AT2G35370 | Glycine cleavage system H protein 1, mitochondrial, GDH1/GCDH, GDCH, GDCSH | N | N | N | Y |
| 135 | | AT3G10920 | Superoxide dismutase [Mn] 1, mitochondrial MSD1 | N | N | N | Y |
| 136 | | AT5G15210 | Zinc-finger homeodomain protein 8, ZHD8/HB30 | N | N | N | Y |
| 137 | (Miller et al. 2013) | AT4G31800 | WRKY transcription factor 18, WRKY18 | N | N | N | Y |
| 138 | | AT4G28400 | Probable protein phosphatase 2C 58 | N | N | N | Y |
| 139 | | AT1G22280 | phytochrome-associated protein phosphatase type 2C, PAPP2C | N | N | N | Y |
| 140 | | AT5G12150 | Rho GTPase-activating protein 6, ROPGAP6/MXC9.11 | N | N | N | Y |
| 141 | | AT4G03280 | Cytochrome b6-f complex iron-sulfur subunit, chloroplastic, PGR1/PETC | N | N | N | Y |

| | | | | | | |
|---|---|---|---|---|---|---|
| 142 | AT5G23540 | Mov34/MPN/PAD-1 family protein MQM1.19/ 26S proteasome non-ATPase regulatory subunit 14 homolog, RPN11 | N | N | N | Y |
| 143 | AT4G16545 | Heat shock protein HSP20 | N | N | N | Y |
| 144 | AT2G30550 | Alpha/beta-Hydrolases superfamily protein DAD1-Like Lipase 3, DALL3 | N | N | N | Y |
| 145 | AT4G02520 | Glutathione S-transferase F2, GSTF2 | N | N | N | Y |
| 146 | AT4G28990 | RanBP2-type domain-containing protein | N | N | N | Y |
| 147 | AT3G57810 | OVARIAN TUMOR DOMAIN-containing deubiquitinating enzyme 4, OTU4 | N | N | N | Y |
| 148 | AT5G61380 | Two-component response regulator-like APRR1, TOC1/AIP1 | N | N | N | Y |
| 149 | AT4G26510 | Uridine kinase-like protein 4, UKL4/UPT1 | N | N | N | Y |
| 150 | AT4G32260 | H+-transporting ATP synthase-like protein/ PIGMENT DEFECTIVE 334, PDE334/atpG//F10M6.100 | N | N | N | Y |
| 151 | AT1G48350 | 50S ribosomal protein L18, chloroplastic, RPL18/EMB3105 | N | N | N | Y |
| 152 | AT1G14000 | VH1-interacting kinase, VIK | N | N | N | Y |
| 153 | AXX17_AT4G27240 | TIR domain-containing protein AXX17_At4g27240 | N | N | N | Y |
| 154 | AT1G32470 | Glycine cleavage system H protein 3, mitochondrial, GDH3 | N | N | N | Y |

| | | | | | | | |
|---|---|---|---|---|---|---|---|
| 155 | | AT1G20570 | Gamma-tubulin complex component, F5M15.11 | N | N | N | Y |
| 156 | | AT3G61380 | TON1 Recruiting Motif 14/ Phosphatidylinositol N-acetyglucosaminlytransferase subunit P-like protein T20K12.280, TRM14/ F2A19.4 | N | N | N | Y |
| | | AT3G50930 | Protein HYPER-SENSITIVITY-RELATED 4, HSR4/ BSC1 | N | N | N | Y |
| 157 | | AT3G16910 | Acetate/butyrate--CoA ligase AAE7, peroxisomal | N | N | N | Y |
| 158 | (Miller et al. 2013) | AT4G30010 | Uncharacterized protein, F6G3_40 | N | N | N | Y |
| 159 | | AT1G52400 | Beta-D-glucopyranosyl abscisate beta-glucosidase, BGLU18 | Y | Y | Y | Y |
| 160 | | AT1G06130 | Probable hydroxyacylglutathione hydrolase 2, chloroplastic, GLX2-4 | Y | Y | Y | Y |
| 161 | (Miller et al. 2010; 2013) | AT1G75660 | 5'-3' exoribonuclease 3, XRN3 | N | Y | N | Y |
| 162 | | AT2G30970 | Aspartate aminotransferase, mitochondrial ASP1 | N | Y | N | Y |
| 163 | (Miller et al. 2013) | AT5G65410 | Zinc-finger homeodomain protein 1, ZHD1/ HB25 | Y | Y | Y | Y |
| 164 | | AT3G50440 | Methylesterase 10, MES10 | N | Y | N | Y |
| 165 | | AT2G41060 | UBP1-associated protein 2B/ RNA-binding (RRM/RBD/RNP motifs) family protein, UBA2B | N | Y | N | Y |
| 166 | | AT1G31330 | Photosystem I reaction center subunit III, chloroplastic PSAF | N | Y | N | Y |
| 167 | (Miller et al. 2010; 2013) | AT3G54960 | Protein disulfide isomerase-like 1-3 PDIL1-3 | N | Y | N | Y |

| | | | | | | | |
|---|---|---|---|---|---|---|---|
| 168 | | AT2G39730 | Ribulose bisphosphate carboxylase/oxygenase activase, chloroplastic RCA | N | Y | N | Y |
| 169 | | AT4G14710 | 1,2-dihydroxy-3-keto-5-methylthiopentene dioxygenase ARD2 | N | Y | N | Y |
| 170 | | AT5G20630 | Germin-like protein subfamily 3 member 3 GER3 | N | Y | N | Y |
| 171 | | AT3G18035 | Winged-helix DNA-binding transcription factor family protein HON4 | N | Y | N | Y |
| 172 | | ATCG00820 | 30S ribosomal protein S19, chloroplastic, rps19 | N | Y | N | Y |
| 173 | | AT4G00180 | Axial regulator YABBY 3, YAB3 | N | Y | N | Y |
| 174 | | AT5G20980 | 5-methyltetrahydropteroyltriglutamate--homocysteine methyltransferase 3, chloroplastic, MS3 | N | Y | N | Y |
| 175 | | AT5G47500 | Pectinesterase, Probable pectinesterase 68, PME68 | N | Y | N | Y |
| 176 | | AT5G57440 | (DL)-glycerol-3-phosphatase 2 GPP2, GS1 | N | Y | N | Y |
| 177 | (Miller et al. 2010) | AT1G07890 | L-ascorbate peroxidase 1, cytosolic, APX1 | N | Y | N | Y |
| 178 | | AT3G21175 | ZIM-like 1, GATA transcription factor 24, ZML1/ GATA24 | N | Y | N | Y |
| 179 | | AT2G19940 | N-acetyl-gamma-glutamyl-phosphate reductase At2g19940 | N | Y | N | Y |
| 180 | | AT2G41530 | S-formylglutathione hydrolase SFGH | N | Y | N | Y |
| 181 | | AT2G24420 | Uncharacterized protein AT2G24420 | Y | Y | Y | Y |
| 182 | | AT3G62240 | RING-type domain-containing protein At3g62240 | N | Y | N | Y |
| 183 | | AT3G18780 | ACTIN-2, LSR2 | N | Y | N | Y |

| | | | | | | | |
|---|---|---|---|---|---|---|---|
| 184 | | AT1G48625 | F-box/kelch-repeat protein At1g48625 | N | Y | N | Y |
| 185 | | AT5G42330 | Uncharacterized protein At5g42330 | N | Y | N | Y |
| 186 | | AT2G13360 | Serine-glyoxylate aminotransferase, AGT1 | N | Y | N | Y |
| 187 | | AT2G26770 | Stomatal closure-related actin-binding protein 1, SCAB1 | N | Y | N | Y |
| 188 | | AT3G60240 | Eukaryotic translation initiation factor 4G, eIF4G | Y | N | N | Y |
| 189 | | AT5G59570 | Transcription factor BOA | Y | N | N | Y |
| 190 | | AT4G14040 | Selenium-binding protein 2, SBP2 | Y | N | N | Y |
| 191 | | AT3G50340 | Uncharacterized protein, F11C1_180 | Y | N | N | Y |
| 192 | | AT1G14450 | NADH dehydrogenase [ubiquinone] 1 beta subcomplex subunit 3-B | Y | Y | N | Y |
| 193 | | AT5G21105 | Plant L-ascorbate oxidase At5g21105 | Y | Y | N | Y |
| 194 | | AT4G17950 | AT-hook motif nuclear-localized protein 13 AHL13 | Y | Y | N | Y |
| 195 | (Miller et al. 2010; 2013) | AT4G38130 | Histone deacetylase 19 HDA19 | Y | Y | N | Y |
| 196 | | AT2G32160 | S-adenosyl-L-methionine-dependent methyltransferases superfamily protein At2g32160 | Y | Y | N | Y |
| 197 | | AT3G02875 | Peptidase M20/M25/M40 family protein ILR1 | Y | Y | N | Y |
| 198 | | AT4G16566 | Histidine triad nucleotide-binding 4, HINT4 OR Bifunctional adenosine 5'-phosphosulfate phosphorylase/adenylylsulfatase HINT4 | Y | Y | N | Y |
| 199 | | AT5G42390 | Stromal processing peptidase, chloroplastic SPP | Y | Y | N | Y |

| | | | | | | | |
|---|---|---|---|---|---|---|---|
| 200 | (Colignon et al. 2017b) | AT4G14960 | Tubulin alpha chain TUA6, TUBA6 | Y | Y | N | Y |
| 201 | (Miller et al. 2010; 2013) | AT4G19880 | Glutathione S-transferase family protein | Y | Y | N | Y |
| 202 | (Miller et al. 2010; 2013) | AT4G14030 | Selenium-binding protein 1 SBP1, FCAALL.79 | Y | Y | N | Y |
| 203 | (Miller et al. 2010) | AT5G25220 | Homeobox protein knotted-1-like 3 KNAT3 | Y | Y | N | Y |
| 204 | | AT3G56490 | Adenylylsulfatase HINT1, HIT3 | Y | Y | N | Y |
| 205 | | AT3G56860 | UBP1-associated protein 2A, UBA2A | Y | N | Y | Y |
| 206 | | AT3G25840 | Protein kinase superfamily protein, At3g25840 | Y | N | Y | Y |
| 207 | | AT3G18530 | ARM repeat superfamily protein, At3g18530 | Y | N | Y | Y |
| 208 | | AT4G31200 | SWAP (Suppressor-of-White-APricot)/surp RNA-binding domain-containing protein | N | Y | Y | Y |
| 209 | | AT4G10340 | Chlorophyll a-b binding protein CP26, chloroplastic, LHCB5 | N | Y | Y | Y |
| 210 | (Miller et al. 2010) | AT5G28300 | Trihelix transcription factor GTL2 | N | Y | Y | Y |
| 211 | | AT4G36870 | BEL1-like homeodomain protein 2 BLH2 | Y | Y | Y | Y |
| 212 | | AT5G21100 | L-ascorbate oxidase AAO | N | Y | Y | Y |
| 213 | | AT4G37930 | Serine hydroxymethyltransferase 1, mitochondrial, SHM1 | N | Y | Y | Y |
| 214 | | AT1G53230 | Transcription factor TCP3 | N | Y | Y | Y |
| 215 | | AT2G34420 | Chlorophyll a-b binding protein, chloroplastic, Lhb1B2 | N | Y | Y | Y |

| | | | | | | | |
|---|---|---|---|---|---|---|---|
| 216 | | AT3G10760 | HTH myb-type domain-containing protein, T7M13.16 | N | Y | Y | Y |
| 217 | | AT1G26910 | 60S ribosomal protein L10-2, RPL10B | N | Y | Y | Y |
| 218 | | AT3G02150 | Transcription factor TCP13, PTF1 | N | Y | Y | Y |
| 219 | (Colignon et al. 2017a) | AT5G60390 | GTP binding Elongation factor Tu family protein, A4 EF1A | N | Y | Y | Y |
| 220 | (Miller et al. 2010) | AT3G45980 | Histone H2B, HTB9 | N | Y | Y | Y |
| 221 | (Miller et al. 2010; 2013) | AT5G42520 | Protein BASIC PENTACYSTEINE6, BPC6 | N | Y | Y | Y |
| 222 | | ATCG00830 | 50S ribosomal protein L2, chloroplastic, rpl2 | N | Y | Y | Y |
| 223 | | AT2G24090 | 50S ribosomal protein L35, chloroplastic, RPL35 | N | Y | Y | Y |
| 224 | (Miller et al. 2013) | AT1G47490 | Polyadenylate-binding protein RBP47C | N | Y | Y | Y |
| 225 | | AT5G01530 | Chlorophyll a-b binding protein CP29.1, chloroplastic, LHCB4.1 | N | Y | Y | Y |
| 226 | | AT3G62250 | Ubiquitin 5/Ubiquitin-40S ribosomal protein S27a-3, UBQ5 | N | Y | Y | Y |
| 227 | | AXX17_AT3G35590 | Uncharacterized protein AXX17_At3g35590 | N | Y | Y | Y |
| 228 | | AT5G63970 | E3 ubiquitin-protein ligase RGLG3 | N | Y | Y | Y |
| 229 | | AT1G21640 | NAD kinase 2, NADK2 | N | Y | Y | Y |
| 230 | | AT3G10570 | Cytochrome P450, family 77, subfamily A, polypeptide 6, CYP77A6 | N | Y | Y | Y |

| | | | | | | | |
|---|---|---|---|---|---|---|---|
| 231 | (Colignon et al. 2017a) | ATCG00130 | ATP synthase subunit b, chloroplastic, atpF | N | Y | Y | Y |
| 232 | (Miller et al. 2010) | AT2G25280 | AmmeMemoRadiSam system protein B, At2g25280 | N | Y | Y | Y |
| 233 | | AT5G08520 | Transcription factor SRM1 | N | Y | Y | Y |
| 234 | (Miller et al. 2010; 2013) | AT3G58120 | Basic leucine zipper 61, BZIP61 | N | N | Y | Y |
| 235 | | AT2G42380 | Basic leucine zipper 34, BZIP34 | N | N | Y | Y |
| 236 | | AT3G20250 | Pumilio 5, PUM5 | N | N | Y | Y |
| 237 | (Miller et al. 2010; 2013) | AT1G71040 | Multicopper oxidase LPR2 | Y | Y | Y | Y |
| 238 | | AT2G17550 | RB1-inducible coiled-coil protein TRM26 | N | N | Y | Y |
| 239 | (Miller et al. 2010; 2013) | AT1G15730 | PRLI-interacting factor L/ Cobalamin biosynthesis CobW-like protein, F7H2 | N | Y | Y | N |
| 240 | | AT3G46640 | Homeodomain-like superfamily protein PCL1 | N | Y | Y | N |
| 241 | | AT1G15820 | Chlorophyll a-b binding protein, chloroplastic Lhcb6 | N | Y | Y | N |
| 242 | (Miller et al. 2010) | AT3G10030 | AA_kinase domain-containing protein, At3g10030/ Aspartate/glutamate/uridylate kinase family protein, T22K18.15 | N | Y | Y | N |
| 243 | | ATCG00480 | ATP synthase beta and epsilon chain atpB | N | Y | Y | N |
| 244 | | AT1G75940 | Beta-glucosidase 20, BGLU20 | N | Y | Y | N |
| 245 | | AT5G67250 | F-box protein SKIP2, FB4 | N | Y | Y | N |

| | | | | | | | |
|---|---|---|---|---|---|---|---|
| 246 | | AT1G31970 | DEA(D/H)-box RNA helicase family protein, STRS1 | N | Y | Y | N |
| 247 | | AT1G54830 | Nuclear transcription factor Y subunit C-3, NFYC3 | N | Y | Y | N |
| 248 | | AT3G58350 | Restricted TEV movement 3, RTM3 | N | Y | Y | N |
| 249 | | AT1G02065 | Squamosa promoter binding protein-like 8, SPL8 | N | Y | Y | N |
| 250 | (Miller et al. 2010) | AT1G44770 | T12C22.4 | N | Y | Y | N |
| 251 | | AT5G08350 | GEM-like protein 4 | N | Y | Y | N |
| 252 | | AT4G15475 | F-box/LRR-repeat protein 4, FBL4 | N | Y | Y | N |
| 253 | | AT1G56170 | Nuclear transcription factor Y subunit C-2, NFYC2 | Y | Y | Y | N |
| 254 | | AT3G20640 | Transcription factor bHLH123 | Y | Y | Y | N |
| 255 | | AT5G42180 | Peroxidase 64, PER64 | Y | N | Y | N |
| 256 | | AN1_LOCUS8082 | Uncharacterized protein AN1_LOCUS8082 | Y | N | Y | N |
| 257 | | AT1G44608 | T18F15.16 | Y | N | Y | N |
| 258 | | AT1G29400 | Protein MEI2-like 5, ML5 | Y | Y | N | N |
| 259 | | AT5G46780 | VQ domain-containing protein, MZA15.20 | Y | Y | N | N |
| 260 | | AT1G59740 | Protein NRT1/ PTR FAMILY 4.3, NPF4.3 | Y | Y | N | N |
| 261 | | AT3G56360 | T5P19_10 | Y | Y | N | N |

Footnote: Y= Protein identified in the system-sets, N= Protein not identified in the system-sets

**Table S3. Prediction of SUMOylation sites in identified SUMO1-substrates by GPS-SUMO 2.0 online server (http://sumosp.biocuckoo.org/online.php).** This table is attached as a separate excel file.

**Table S4.** Predicted frequency of SUMO sites identifications across different studies and in the respective host proteome.

| Data set | Number of proteins identified | Number of predicted SUMO sites | Number of proteins without SUMO site | Average SUMO site/protein | % proteins with SUMO sites |
|---|---|---|---|---|---|
| This study* | 261 | 766 sites in 221 proteins | 40 | 2.93 | 84.67 |
| AtSUMO1 SUMOylome during heat shock (Miller et al. 2010) | 357 | NA | NA | 2.24 | 80 |
| Arabidopsis proteome (Miller et al. 2010) | 27,379 | NA | NA | 0.84 | 47 |
| *Solanum tuberosum* leaf SUMOylome (Colignon et al. 2017a) | 77 | 50 sites in 39 proteins | 38 | 0.64 | 50.64 |
| HsSUMO2 SUMOylome (Golebiowski et al. 2009) | 759 | 1681 | 195 | 2.2 | 74.30 |
| Human proteome (Golebiowski et al. 2009) | 43,964 | 28,078 | 28,571 | 0.6 | NA |

Footnote: *SUMO site prediction was performed using GPS-SUMO 2.0 online server (http://sumosp.biocuckoo.org/online.php) (Zhao et al. 2014)

NA: Not reported by authors

**Table S5. Gene ontology (GO) enrichments in 261 SUMO1-substrates identified in this study performed by STRING database (https://string-db.org).** This table is attached as a separate excel file.

**Table S6: List of primers used in this study**

| No. | Primer Name | Forward primer sequence (5’ to 3’) | Reverse primer sequence (5’ to 3’) |
|---|---|---|---|
| 1 | *SAND* RT | AACTCTATGCAGCATTTGATCCACT | TGATTGCATATCTTTATCGCCATC |
| 2 | *PR1* RT | GTGGGTTAGCGAGAAGGCTA | ACCTTGGCACATCCGAGTCT |
| 3 | *PR2* RT | TCAAGGAAGGTTCAGGGATG | TTCACGAGCAAGGGAGATTG |
| 4 | *ICS1* RT | TACTAACCAGTCCGAAAGACG | GAGGCTTGACAACAACTCTGT |
| 5 | *SUMO1* RT | CTTGTTTGATGGGCGTCGTC | CAGTCTGATGGAGCATCGCA |
| 6 | *SUMO2* RT | TCGATGCAATGCTTCATCAGAC | ACCGCCACTAAAAGCAGAAGA |
| 7 | *SUMO3* RT | CGAGCAAATCAGCGTCAGTG | ACACACACGATAACCGACCA |
| 8 | *SAE1* RT | ATTCCTCGGAGAACAGCAAA | ACGCCCTTCACTCTCTTCAA |
| 9 | *SAE2* RT | ACAGCCTTTTTGAAGCGAAA | ATAATGGCGTTGGTCGTAGC |
| 10 | *SCE1* RT | AATGGTGTGGCATTGCACTA | TCCTCACTGAAGTGCATCGT |
| 11 | *SIZ1* RT | AACAGGGAAAGAAGCAGGAA | GGCAGCTTGTTCATCAGAAA |
| 12 | *HPY2* RT | CTACACCTTCCTCAGTGCCA | AACATTCCAAACTGCTTCCC |
| 13 | *ESD4* RT | TGATGCTGGATTCGTTGTCG | TCTCCGCACTTTGCATAAGC |
| 14 | *ELS1* RT | TGGATGGTTACCACAAACGG | ATGAACTTGTCTCCGCAACG |
| 15 | *His-H89R SUM1* (Genotyping) | CAAGCTTGCATGCCTGCAGGTCG | TAGACGTCCGTGAAAGGTTCTCCGTCC |
| 16 | *srfr1-4* (Genotyping) | TCTCCACTGTACTAATTTCCCT | ACTAATTCCGCAACGTGCCT (37460-R) |
| 17 | *sum1-1* (Genotyping) | TTTCGTGTAGCTGCGATTAGG | TTATCTTTGCTCGCCATTAGC |
| 18 | LB3 (Genotyping) | TAGCATCTGAATTTCATAACCAATCTCGATACAC | |
| 19 | HA-SRFR1g specific | ATGTATCCTTATGATGTTCCAGATTATGCT | |
